# Genomic studies in *Linum* shed light on the evolution of the distyly supergene and the molecular basis of convergent floral evolution

**DOI:** 10.1101/2025.04.11.648331

**Authors:** Panagiotis-Ioannis Zervakis, Zoé Postel, Aleksandra Losvik, Marco Fracassetti, Lucile Solér, Estelle Proux-Wéra, Ignas Bunikis, Allison Churcher, Tanja Slotte

## Abstract

- Distyly, an example of convergent evolution, is governed by a supergene called the *S-* locus. Recent studies highlight similar genetic architectures of independently evolved *S*-loci, but whether similar regulatory pathways underlie convergent evolution of distyly remains unclear.
- We examined the evolution of supergenes and mechanisms underlying distyly in *Linum* species that diverged ∼33 Mya. Using haplotype-resolved genomes and population genomics, we identified and characterized the *S-*loci of *Linum perenne* (distylous) and *Linum grandiflorum* (style length dimorphic), and compared them to that of *Linum tenue* (distylous). We then tested for a conserved hormonal mechanism regulating style length polymorphism in *Linum*.
- Hemizygosity in short-styled individuals is a shared feature of the *Linum S*-locus supergene, though its size, gene content, repeat elements, and extent of recombination suppression vary greatly among species. Two distyly candidate genes, *TSS1* (style length) and *WDR-44* (anther height/pollen self-incompatibility) are conserved at the
- *S-*locus. Consistent with a brassinosteroid-dependent role of *TSS1*, epibrassinolide treatment revealed a conserved, morph-specific effect on style length.
- *S-*locus genetic architecture, key *S-*locus genes and mechanisms regulating style length remain conserved >30 Mya in *Linum*. In combination with findings from other systems, our results suggest that the brassinosteroid pathway frequently contributes to style length polymorphism.

## Introduction

Distyly is a floral polymorphism that promotes outcrossing and has evolved repeatedly in flowering plants, making it a prominent example of convergent floral evolution (Barrett, 2019). In distylous species, there are two floral morphs that differ reciprocally in the positions of anthers and stigmas. Long-styled (L-morph, or pin) plants have stigmas in a high position in the flower, and anthers in a low position, whereas short-styled (S-morph, or thrum) individuals have the opposite arrangement. Differences in flower structure are usually accompanied by differences in pollen and stigma traits and by heteromorphic self-incompatibility (SI), which limits self- and intra-morph pollination. Distyly has evolved independently multiple times in flowering plants (Lloyd & Webb, 1992; Naiki, 2012) suggesting that it provides a solution to a common set of selective pressures (Shore *et al*., 2019; Simón-Porcar *et al*., 2024). Specifically, reciprocal morph differences increase the precision of pollen transfer by pollinators (Darwin, 1877; Lloyd & Webb, 1992; Barrett, 2019; Simón-Porcar *et al*., 2024), whereas heteromorphic SI confers inbreeding avoidance (Charlesworth & Charlesworth, 1979).

In distylous species where genetic studies have been done, distyly is governed by a single Mendelian locus, the *S-*locus, with one dominant and one recessive allele (Bateson & Gregory, 1905; Laibach, 1923; reviewed by Ganders, 1979) which controls both floral morphology and heteromorphic SI. In most systems, the L-morph is homozygous for the recessive *s-*allele (*s/s*), whereas the S-morph is genetically heterozygous (*S/s*) (reviewed by Ganders, 1979). To explain how a single Mendelian locus could control this multi-trait balanced polymorphism, Ernst (1936) proposed that in distylous *Primula*, the *S-*locus harbored at least three separate and polymorphic genes, present in close linkage and controlling different aspects of distyly. Under Ernst’s model, the *S*-locus constitutes a supergene, defined as “a system of closely linked loci controlling a polymorphic phenotype, such that a non-recombining genome region is structured into two or more distinct haplotypes, each carrying a set of alleles that control multiple aspects of one of the phenotypes” (Charlesworth, 2016).

Detailed genomic characterization of multiple independently evolved distyly *S-*loci has shown that they harbor multiple closely linked genes important for trait polymorphism, and can be considered supergenes (e.g., *Fagopyrum* (Yasui *et al*., 2012; Fawcett *et al*., 2023); *Primula* (Huu *et al*., 2016; Li *et al*., 2016); *Turnera* (Shore *et al*., 2019); *Linum* (Gutiérrez-Valencia *et al*., 2022); *Nymphoides* (Yang *et al*., 2023); *Gelsemium* (Zhao *et al*., 2023); Oleaceae (Castric *et al*., 2024; Raimondeau *et al*., 2024). However, unlike other types of supergenes, which often harbor inversions, all distyly supergenes studied in detail so far instead harbor large indels (reviewed in Gutiérrez-Valencia *et al*., 2021b). Because the dominant allele is typically longer and the recessive allele shorter, the supergene is usually hemizygous in S-morph individuals, with S-morph-specific expression of dominant *S-*linked genes that control floral morph (reviewed in (Gutiérrez-Valencia *et al*., 2021b)). Hemizygosity ensures both dominant expression and absence of recombination between the recessive and dominant alleles at the distyly supergene.

Whether repeated evolution of similar genomic architectures at the distyly supergene is accompanied by functional similarities in the mechanisms regulating distyly remains less clear, as independently evolved *S-*loci do not share orthologous genes. Genomic studies have enabled experimental and functional studies of candidate *S-*locus genes and mechanisms governing distyly, demonstrating that *S-*linked genes that inactivate brassinosteroids control style length and female SI in at least two distylous systems (in *Primula* (Huu *et al*., 2016, 2022) and in *Turnera* (Shore *et al*., 2019; Matzke *et al*., 2020, 2021)). Studies in additional distylous systems are required to determine whether distyly is generally accompanied by parallel evolution at the biochemical pathway level.

*Linum* (wild flaxseed species) is a classic system for the study of the function, evolution and genetic basis of distyly (Darwin, 1863, 1877; Dulberger, 1992; Armbruster *et al*., 2006). This system is of particular interest due to its varied stylar polymorphisms, including distyly (Fig. 1b-c) and stigma height dimorphism (Fig. 1d) (Armbruster *et al*., 2006; McDill *et al*., 2009; Ruiz-Martin *et al*., 2018; Maguilla *et al*., 2021). The presence of varied stylar polymorphisms, as well as recurrent loss of distyly, make *Linum* a particularly suitable system for dissecting the genetic basis of distyly.

**Fig. 1.**
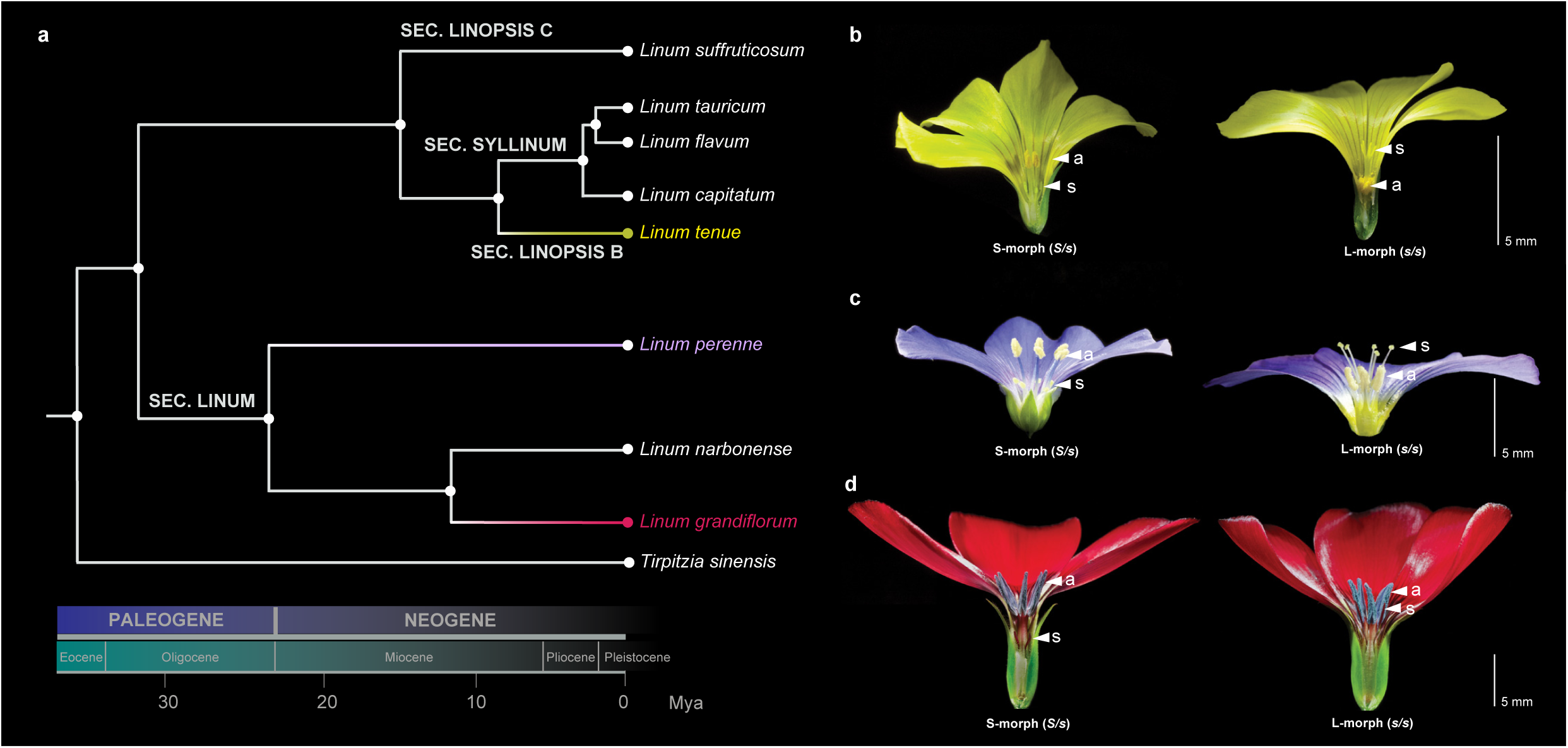
Schematic phylogeny and floral morphs of the study species. **a**. Schematic phylogeny and divergence times of the species used in the study. The three main species of interest (*L. grandiflorum, L. perenne* and *L. tenue*) are highlighted with color. **b.** Floral morph of S-morph (left) and L-morph (right) *L. tenue*. **c**. Floral morph of S-morph (left) and L-morph (right) *L. perenne*. **d.** Floral morph of S-morph (left) and L-morph (right) *L. grandiflorum*. In panels **b-d**, Positions of anthers (a) and stigmas (s) are marked in the Fig. and magnification is indicated by a scale bar (5 mm). Part of the corolla and sepals were removed for improved visibility of sexual organ location.

Building on a high-quality genome assembly of the distylous and SI *Linum tenue* (Fig. 1b), we recently characterized the distyly *S-*locus in *Linum*, and showed that in *L. tenue* it constitutes a supergene which harbors an ∼260 kb indel, rendering the S-morph predominantly hemizygous (Gutiérrez-Valencia *et al*., 2022). The *L. tenue S-*locus harbors nine protein-coding genes, including candidate genes for style length (*LtTSS1*) and anther height/pollen SI (*LtWDR-44*). In the closely related selfing species *L. trigynum* which recently lost distyly and is homostylous, i.e., monomorphic with anthers and stigmas at the same height, *LtWDR-44* is present but expressed a lower level than in SI *L. tenue* thrums (Gutiérrez-Valencia *et al*., 2024). Altered expression of *LtWDR-44* is associated with a switch in pollen SI function from thrum-to-pin-type and self-compatibility (SC), suggesting a role for this gene in pollen SI (Gutiérrez-Valencia *et al*., 2024).

Based on our previous work in *L. tenue* and *L. trigynum* (Gutiérrez-Valencia et al. 2022; 2024), two genes thus stand out as strong candidate genes for governing distyly in *Linum*: *LtTSS1* (hereafter called *TSS1*) and *LtWDR-44* (hereafter called *WDR-44*). However, it is not currently clear whether these two genes are generally important for distyly in *Linum,* as ancestral state reconstruction suggested that divergent *Linum* species may have independently evolved distyly (Armbruster *et al*., 2006; McDill *et al*., 2009; Ruiz-Martin *et al*., 2018). To shed further light on the origin(s) of the distyly *S-*locus in *Linum*, and to identify conserved candidate genes and mechanisms regulating distyly in *Linum*, there is a need for further comparative studies of the *S-*locus, building on high-quality genome assemblies of widely diverged distylous *Linum* species.

Here, we describe new high-quality haplotype-resolved genome assemblies of two *Linum* species: the distylous and SI *L. perenne* and the style length dimorphic and SI *L. grandiflorum*, both of which diverged from *L. tenue* about 33 Mya (Maguilla *et al*., 2021) (Fig. 1a). Like *L. tenue,* both *L. perenne* and *L. grandiflorum* have heteromorphic SI (Murray, 1986), but *L. grandiflorum* lacks anther height dimorphism. To identify shared *S-*locus genes that might be involved in the control of floral morph and SI in *Linum*, we 1) identify and characterize the *S-*loci of *L. grandiflorum* and *L. perenne* and compare their gene content to that of *L. tenue*. We use the results to 2) infer the origin of the distyly *S-*locus in *Linum*, and 3) experimentally test whether downregulation of brassinosteroid-responsive genes by the distyly supergene is a conserved mechanism that controls style length polymorphism in *Linum*. Our results shed new light on the origin and evolution of the distyly supergene in *Linum* and the genetic basis of an iconic case of convergent floral evolution.

## Materials and Methods

### Biological material for genome assembly and annotation

For *de novo* genome assembly of *L. perenne* L. and *L. grandiflorum* Desf., we snap-froze leaves from one S-morph individual of *L. perenne* accession LIN 2003 (here named L96A) and one S-morph individual of *L. grandiflorum* accession LIN 10 (here named L62.06), both from the IPK Gatersleben Gene Bank (Table S1). For annotation of genome-assemblies, we snap-froze at least two replicates each of leaves, stems, early and late flower buds (collected at two stages for *L. perenne* and at three stages for *L. grandiflorum*) and open flowers of *L. perenne* L96A and *L. grandiflorum* L62.06 for RNA extraction and sequencing.

### Plant growth conditions

Seeds were sterilized with 10% bleach or 3% hydrogen peroxide and detergent, followed by washes with 70% ethanol and distilled water, sown on sterile plates with half strength Murashige-Skoog medium (Sigma-Aldrich, St. Louis, MO, USA) stratified and moved to standard long-day conditions at 16 h light at 20°C: 8 h dark at 18°C, 60% maximum humidity, 120 μE light intensity, until seedlings emerged. Seedlings were transplanted to pots containing a mixture of soil (Hasselfors Garden, Sweden) and gravel (1.5:1), with an addition of perlite and vermiculite. *L. perenne* plants were vernalized for 9 weeks under short-day conditions: 8 h light at 6°C, 16 h dark at 2°C, 65% maximum humidity, 110 μE light intensity, with the following transition conditions were in place two weeks before and after vernalization: 11 h light at 15°C, 13 h dark at 10°C, 120 μE light intensity, 65% maximum humidity.

### High Molecular Weight (HMW) DNA isolation

HMW DNA was extracted and purified using a two-step protocol, from a total of 2 g fresh-frozen leaves from individuals *L. perenne* L96A and *L. grandiflorum* L62.06. For HMW DNA extraction we followed a modified protocol from (Fulton *et al*., 1995) with purification using Genomic-Tip/500 (Qiagen, Hilden, Germany). HMW DNA quality was checked spectrophotometrically and through pulsed-field gel electrophoresis using SeaKem Gold agarose (Lonza, Rockland, ME, USA), 0.5X KBB buffer (Sage Science), and a Pippin Pulse Electrophoresis power supply system (Sage Science, Beverly, MA, USA), with post-staining using GelRed (Biotium, Fremont, CA, USA).

### PacBio High-Fidelity (HiFi) sequencing

HMW DNA from *L. perenne* L96A (32 μg) and *L. grandiflorum* L62.06 (33 μg) was used to generate SMRTBell libraries for HiFi long-read sequencing. Each library was sequenced on two SMRT cells in HiFi mode on a Sequel II (Pacific Biosciences), which resulted in 31 and 50 Gbases of HiFi data for *L. perenne* and *L. grandiflorum*, respectively, with an insert size of 15 kbp.

### Hi-C data generation

To generate high-quality proximity ligation libraries (Hi-C) for scaffolding of genome assemblies, a total of 300 mg of fresh-frozen leaf tissue was first ground to a fine powder in liquid nitrogen. Hi-C libraries were generated using the Dovetail OmniC kit. Sequencing on an Illumina NovaSeq6000 generated a total of 1.0*10^9^ paired-end 150 bp reads for *L. perenne*, and 2.3*10^9^ paired-end 150 bp reads for *L. grandiflorum*.

### RNA extraction and sequencing

For genome annotation, we obtained RNA sequencing data from leaves, stems, flower buds and open flowers of *L. perenne* L96A and *L. grandiflorum* L62.06. Total RNA was extracted using the RNeasy Plant Mini Kit (Qiagen, Hilden, Germany). Sequencing libraries were prepared using the TruSeq stranded mRNA library preparation kit (Illumina, San Diego, CA, USA) including polyA selection and unique dual indexes (Illumina, San Diego, CA, USA), and were sequenced using paired-end 150bp reads on a NovaSeq6000 system.

### De novo genome assembly

We generated primary and haplotype-resolved genome assemblies based on HiFi and Hi-C data of our outbred S-morph *L. perenne* L96A and *L. grandiflorum* L62.06 individuals using integrated Hi-C assembly settings in Hifiasm (Cheng *et al*., 2021). For each species, we generated two high-quality phased haplotype assemblies, designated as hap1 and hap2, as well as a primary assembly. Assembly completeness was checked using Benchmarking Universal Single-Copy Orthologs (BUSCO) (Waterhouse *et al*., 2018) against the eudicots_odb10 gene data set. Prior to annotation, assemblies were screened for contamination (Note S1) and presence of chloroplast or mitochondrial sequences as described in (Gutiérrez-Valencia *et al*., 2024).

### Genome annotation

Annotation of genes and repeats was performed using open-source pipelines in use at the National Bioinformatics Infrastructure Sweden (NBIS) Annotation and Assembly unit (See Data Availability). We used a combination of evidence-based and *ab initio* annotation, followed by functional annotation. In addition, repeats were modeled and annotated, after vetting them against annotated genes. We fully annotated the primary and haplotype-resolved assemblies of each species.

For evidence-based gene annotation methods we used both proteins and transcriptomes. As protein evidence, we used proteins from sequenced *Linum* species (*L. tenue, Linum usitatissimum*), more distantly related species from the Malpighiales (*Manihot esculenta, Populus trichocarpa, Ricinus communis, Salix purpurea*), the Vitales (*Vitis vinifera*) as well as Uniprot data for rosids. We further used transcriptome data from leaves, stems, buds and flowers of *L. perenne* L96A and *L. grandiflorum* L62.06. After adapter-trimming with fastp v0.23.2 (Chen *et al*., 2018), RNAseq reads were aligned to the reference genome using Hisat2 v2.1.0 (Kim *et al*., 2015). Genome-guided assembly of transcripts was done using StringTie v2.2.1 (Pertea *et al*., 2015), using MultiQC (Ewels *et al*., 2016) for quality-checking. Evidence-based annotation was performed using MAKER v3.01.02 (Holt & Yandell, 2011) including aligned transcript sequences and reference proteins as evidence, whereas *ab-initio* training was conducted using GeneMark v4.3 (Besemer *et al*., 2001), Augustus v3.3.3 (Stanke *et al*., 2006), and Snap 2013_11_29 (Korf, 2004). Finally, results from *ab-initio* and evidence-based annotation were combined to produce final gene builds, which were functionally annotated using Blast (v. 2.9.0) (Altschul *et al*., 1990) matches against Uniprot/Swissprot and results from InterproScan 5.59-91.0 (Hunter *et al*., 2012).

Species-specific repeat libraries were generated using RepeatModeler (Smit & Hubley, 2008), and candidate repeats were vetted against protein evidence (excluding transposons) to exclude low-complexity coding sequences. Finally, repeat identification was performed using RepeatMasker (Smit *et al*., 2013) and RepeatRunner (Smith *et al*., 2007).

### Manual curation of annotation in the S-locus region

To describe and compare the gene content of the *S-*locus region across species, we manually curated gene annotation in genome regions of *L. perenne* and *L. grandiflorum* containing their respective S-morph hemizygous *S*-locus (*L. perenne* h1tg000002l:1,080,000-4,890,000; *L. grandiflorum* h1tg000023l: 11,240,000-12,420,000 – see section “Identification of the *S-* locus in *L. perenne* and *L. grandiflorum*” below for details) by inspecting transcriptome evidence for the original annotation as well as for TransDecoder/StringTie v2.2.1 based gene predictions. Manual curation resulted in removal of eight gene models and addition of two gene models in the *L. perenne* hemizygous *S*-locus region, and removal of 14 gene models that were not supported by transcriptome data, and addition of five new gene models based on TransDecoder (https://github.com/TransDecoder/TransDecoder) output in the *L. grandiflorum* hemizygous *S-*locus region.

We performed additional transposable element (TE) and repeat annotation to improve TE classification completeness prior to tests for repeat enrichment at the *S-*locus. Specifically, we used HiTE v3.2 (Hu *et al*., 2024) in conjunction with LTR_retriever v2.9.9 (Ou & Jiang, 2018) to build a repeat element library and annotate the genome with full-length TEs, classified using RepeatMasker v4.1.5 (Smit *et al*., 2013) in sensitive mode. Statistical comparison of *S-*locus and genome-wide repeat content was done using binomial tests in R v4.3.2.

### Whole-genome short-read sequencing

DNA for short-read sequencing was extracted from 157 samples of *L. perenne* from three natural populations and two accessions, and for *L. grandiflorum* we extracted DNA from 22 individuals from three accessions (Table S1), using the *Quick*-DNA Miniprep Plus Kit (Zymo Research, Irvine, CA, USA). We also acquired short-read sequencing data for five additional distylous species of *Linum* (Fig. 1; Table S1) following the same procedure but using magnetic beads and the Quick-DNA MagBead Plus kit (Zymo Research, Irvine, CA, USA) for DNA extraction. Sequencing libraries were prepared from 1 μg DNA using the TruSeq PCRfree DNA sample preparation kit (Illumina, San Diego, CA, USA) with unique dual indexes, targeting an insert size of 350 bp. Libraries were sequenced on a NovaSeq 6000 system, yielding paired-end 150 bp reads.

### Short-read processing, mapping, variant calling and filtering

Illumina whole-genome resequencing reads were adapter- and quality-trimmed using BBDuk from BBMap v38.61b (Bushnell, 2014), and mapped using BWA-MEM v0.7.18 (Li, 2013). We excluded mapped reads with a mapping quality lower than 20 and duplicates using Picard tools v3.1.1 (http://broadinstitute.github.io/picard). Variants were called using BCFtools *mpileup* v1.17 (Danecek & McCarthy, 2017) independently for each species. We kept only bi-allelic variants and invariant sites, and applied additional filters for depth, missingness and mapping quality (BCFtools *min_depth* = 5; *max_depth* = 200; *missingness* = 0.9; *min_quality* = 20). Due to the high repeat content of our assemblies additional masking of repeats was necessary. Hence, we masked repeats using ‘*bedtools intersect*’ and by filtering on coverage as in (Gutiérrez-Valencia *et al*., 2021a). Finally, to reduce false heterozygous calls, we applied an allele balance filter with thresholds 0.2 and 0.8, setting heterozygous calls that failed this criterion to missing.

### Identification of the S-locus in L. perenne and L. grandiflorum

To identify the *S-*locus we tested for an association between single nucleotide polymorphism (SNP) genotype and floral morph using genome-wide association mapping (GWAS). Prior to GWAS we removed sites with missing data, rare variants (minor allele frequency < 0.05), and pruned variants with high linkage disequilibrium (LD) (*r^2^* > 0.2) in 50 kb windows. We performed association analysis in PLINK v1.90b4.9 (Purcell *et al*., 2007) using Fisher’s exact test on genotypes, assuming a dominant effect for the minor allele and applying a False Discovery Rate (FDR) P-value adjustment. In *L. grandiflorum,* this association analysis used 15,927 LD pruned SNPs in 11 S-morph and 7 L-morph individuals from three accessions (Table S1). In *L. perenne*, we first analyzed 13,992 LD-pruned SNPs genome-wide from 7 S-morph and 12 L-morph *L. perenne* individuals from one family. Because family-based analyses can have limited resolution, we validated our findings by GWAS analyses on 53 individuals from one natural population (ger3, Table S1; Note S2).

We performed depth of coverage analyses to identify genomic regions with presence-absence variation between morphs in the genomes of *L. perenne* and *L. grandiflorum* and narrow down the position of the *S-*locus. Depth of coverage of reads mapped to the hap1 haplotype-resolved assembly of each species was calculated in 300kb kb windows using BEDTools v2.31.1 (Quinlan & Hall, 2010) and normalized by total sample read count. We identified windows that differed in normalized median coverage between individuals with different floral morph (L-vs S-morph) using a two-sample Fisher-Pitman permutation test in R (v.4.3.2, package “coin” v1.4.3), with 1,000,000 permutations, using a significance threshold of (P*≤0.01*) after Bonferroni multiple testing correction.

### Stepwise assembly of the S-locus gene set

We estimated *d_S_* between *S-*locus genes and their closest paralogs in *L. grandiflorum* and *L. perenne*, to test for an impact of stepwise gene movement on *S-*locus gene content. To do so, we estimated *d_S_* between each *S*-locus gene and its closest paralog, identified by OrthoFinder v2.5.5 (Emms & Kelly, 2019). We estimated *d_S_* in MEGA X (Tamura et al., 2021) under the Nei-Gojobori model. Widely different *d_S_* estimates for different genes imply stepwise gene duplication at different times, and very low estimates suggest very recent gene duplication.

### Selection on S-locus candidate genes and the age of the S-locus in Linum

To assess purifying selection on the *S-*locus candidate genes *TSS1* and *WDR-44*, we estimated the ratio of nonsynonymous to synonymous sequence divergence (*d_N_/d_S_*) for both genes across *L. perenne, L. grandiflorum*, *L. tenue* and five additional distylous species of *Linum* (Fig. 1; Table S1). We obtained *TSS1* and *WDR-44* sequences of the additional distylous *Linum* using the HybPiper pipeline (Johnson *et al*., 2016), based on a target file including available gene sequences of *TSS1* and *WDR-44* from *L. tenue* and *L. trigynum* (Gutiérrez-Valencia *et al*., 2022, 2024), *L. grandiflorum* and *L. perenne*. We constructed multiple sequence alignments for coding sequences of each gene using T-Coffee (Notredame *et al*., 2000), inferred gene trees using RAxML with the GTR-Gamma model of nucleotide substitution (Stamatakis, 2014) and estimated *d_N_/d_S_* ratios using codeml in PAML (Yang, 2007), comparing a model allowing different values of *d_N_/d_S_* for each branch to a constrained model with only one *d_N_/d_S_* ratio for the whole tree using a likelihood ratio test (LRT).

Next, we took advantage of the presence of *WDR-44* paralogs to compare selective pressures on the *S*-locus sequence compared to its paralogs and to infer the timing of duplication. To compare selective pressures on the *S-*locus vs paralogs of *WDR-44* we used the codeml branch model and three phylogenetic tree annotations: (i) only one *d_N_/d_S_* ratio for the whole tree, (ii) one *d_N_/d_S_* ratio for *S*-locus sequences and one for paralogs, and (iii) one *d_N_/d_S_* ratio per species, with model selection based on LRTs. To estimate the timing of duplication of *WDR-44*, we ran BEAST2 v2.7.7 (Bouckaert *et al*., 2019) with a lognormal optimized relaxed molecular clock and a calibrated Yule model. For calibration we used the timing of diversification of *Linum* species, i.e., ∼33 Mya (parameters: lognormal, M = 3.5, S = 0.05) (Maguilla *et al*., 2021). We set the chain length to 100,000,000, sampling every 10,000th step and obtained the final estimate after excluding the first 10% of trees as burn-in. We also applied a simple molecular clock analysis to estimate the divergence time of *L. tenue* and *L. perenne* (*t = d_S_/(2μ)*), based on synonymous sequence divergence at *TSS1* and *WDR-44* estimated under the Nei-Gojobori model in MEGA X and assuming a mutation rate (*μ)* of 7*10^−9^ (Ossowski et al. 2010). Finally, we included *WDR-44* sequences for three outgroups in our phylogenetic reconstruction. For *Tirpitzia sinensis*, the *WDR-44* sequence was retrieved using the same procedure as for the *Linum* species. For *Manihot esculenta* we used *Manes.15G085300* for *M. esculenta* and for *Populus trichocarpa Potri.011G122500* as in (Gutiérrez-Valencia *et al*., 2022).

### Brassinosteroid supplementation experiment

To test whether supplementation with active brassinosteroid hormone affects style length specifically in the S-morph of both *L. perenne* and *L. tenue*, we performed a controlled experiment with two treatments: eBL treatment (10 μM 24-epibrassinolide (eBL) dissolved in 0.1% dimethylsulfoxide, DMSO) and control treatment (0.1% DMSO). eBL concentration was chosen based on initial tests with 1 μM, 10 μM, and 20 μM eBL. Young flower buds were injected until saturation on two consecutive days with with either eBL or control solution. Fully open flowers were dissected, photographed under a stereo microscope (Leica S APO) and style and stamen lengths were determined using ImageJ 1.53k (Schneider *et al*., 2012). One to two flowers of eight L-morph and 11 *L. perenne* S-morph individuals were subjected to each treatment type (eBL or control) for a total of 90 measurements of style and stamen length. For *L. tenue*, two flowers of each of 19 L-morph and S-morph individuals were subjected to each treatment type, for a total of 152 measurements of style and stamen length. The experiment was not performed on *L. grandiflorum* due to growth chamber space and time constraints.

Style and stamen lengths were analyzed separately using analysis of variance (ANOVA) in R v4.1.1 using the lm() function, with organ length as the response and floral morph, hormonal treatment, and the interaction between floral morph and treatment as predictors. For models with significant effects, we conducted a post-hoc test using the Tukey “Honest Significant Difference” (HSD) method and obtained 95% CIs for the difference in mean organ length.

To test whether eBL treatment affects style cell length, we quantified cell length in styles subjected to eBL or control treatment in *L. perenne*. To obtain an image of the epidermal cells in control and treated styles, a thin layer of UV-cured transparent nail polish (Semilac, Poznan, Poland) was applied to a microscope slide and excised styles were carefully placed on the surface of the nail polish (this method was not feasible for fixed *L. tenue* material and therefore it was only performed in *L. perenne*). After hardening under the UV-light the imprint was photographed under a light microscope (Olympus BX60) and cell sizes were measured using ImageJ 1.53k (Schneider *et al*., 2012). Cell length measurements were obtained separately for three different sections at the bottom, middle and top of the style (10 cells measured per section), following (Ushijima *et al*., 2015) and (Foroozani *et al*., 2023). Measurements were performed on two to four flowers of each of eight *L. perenne* L-morph individuals and 11 S-morph individuals, resulting in a total of 805 cell length measurements.

The 10 cell length measurements for each style section were averaged prior to linear model analysis using lm() in R v4.1.1. We tested for an effect of eBL treatment on mean style cell length using a linear model with mean cell length as the response, and floral morph, style section (bottom, middle, or top) and hormonal treatment as predictor variables. Cell lengths were log-transformed to improve normality of residuals. Post-hoc tests were performed and 95% CI intervals obtained for significant effects as described above.

## Results

### High-quality phased genome assemblies of Linum perenne and Linum grandiflorum

As both *L. grandiflorum* and *L. perenne* are SI (Murray 1986) and outbred, we assembled both a primary assembly and a pair of haplotype-resolved assemblies for each species based on PacBio HiFi and Hi-C data. We sequenced S-morph individuals which are expected to harbor both the dominant and recessive alleles at their *S-*loci. Hi-C integrated assembly in Hifiasm (Cheng *et al*., 2021) resulted in highly complete primary and haplotype-resolved assemblies with BUSCO scores ranging from 94.0% to 95.1% (Table S2) that were highly contiguous, with N50 scores ranging from 10.2 Mb to 69.9 Mb (Table S2). Assembly lengths were similar to genome sizes estimated by flow cytometry (Table S2).

We annotated the primary and haplotype-resolved assemblies using a combination of *ab-initio* and evidence-based methods. We identified a total of ∼42,000 protein-coding genes in our *L. grandiflorum* assemblies, whereas our *L. perenne* assemblies had ∼45,000 protein-coding genes (Data S1A). Compared to *L. tenue*, where 49.4% of the genome consisted of repeats (Gutiérrez-Valencia *et al*., 2022), the genomes of *L. grandiflorum* and *L. perenne* were richer in repeats, with 78.2% and 69.5% of the respective genome assemblies annotated as repetitive (Data S1A). The relatively high gene numbers of *L. perenne* and *L. grandiflorum* likely result from an ancient whole genome duplication in the ancestor of these species (Sveinsson *et al*., 2014).

### Hemizygosity in the S-morph is a common feature of Linum S-locus supergenes

To identify *S*-loci in *L. perenne* and *L. grandiflorum* we searched for single nucleotide polymorphisms (SNPs) whose genotypes were associated with floral morph. Because many distyly *S*-locus supergenes harbor presence-absence variation, we also tested for presence-absence variation between floral morphs using short-read depth of coverage analyses.

In *L. grandiflorum,* GWAS identified two SNPs on contig h1tg000023l of haplotype-resolved assembly hap1 as significantly associated with floral morph (Fisher exact test, assuming dominant effect of the S-morph-specific allele, FDR<0.05) (Figs 2a, S1). The associated SNPs define an ∼1.2 Mb region on contig h1tg000023l ranging from ∼11.2 Mb to ∼12.4 Mb. Within this region, coverage analyses showed presence-absence variation between floral morphs, with significantly lower normalized median coverage in L-morph than S-morph individuals (median normalized coverage 0 for L-morph and 14.8 for S-morph, permutation test with 1,000,000 resamples, P<0.01 after Bonferroni correction, Fig. 2a). These results suggest that h1tg000023l on haplotype-resolved assembly hap1 harbors the longer, dominant allele at the *L. grandiflorum S-*locus. Comparison between the two haplotype-resolved assemblies of *L. grandiflorum* confirmed the presence of a ∼1.2 Mb hemizygous region in S-morph individuals and identified h2tg000012l on the hap2 assembly as harboring the shorter, recessive *S-*allele. Although the recessive allele was shorter than the dominant allele, it included a unique 70 kb region missing from the dominant allele.

**Fig. 2.**
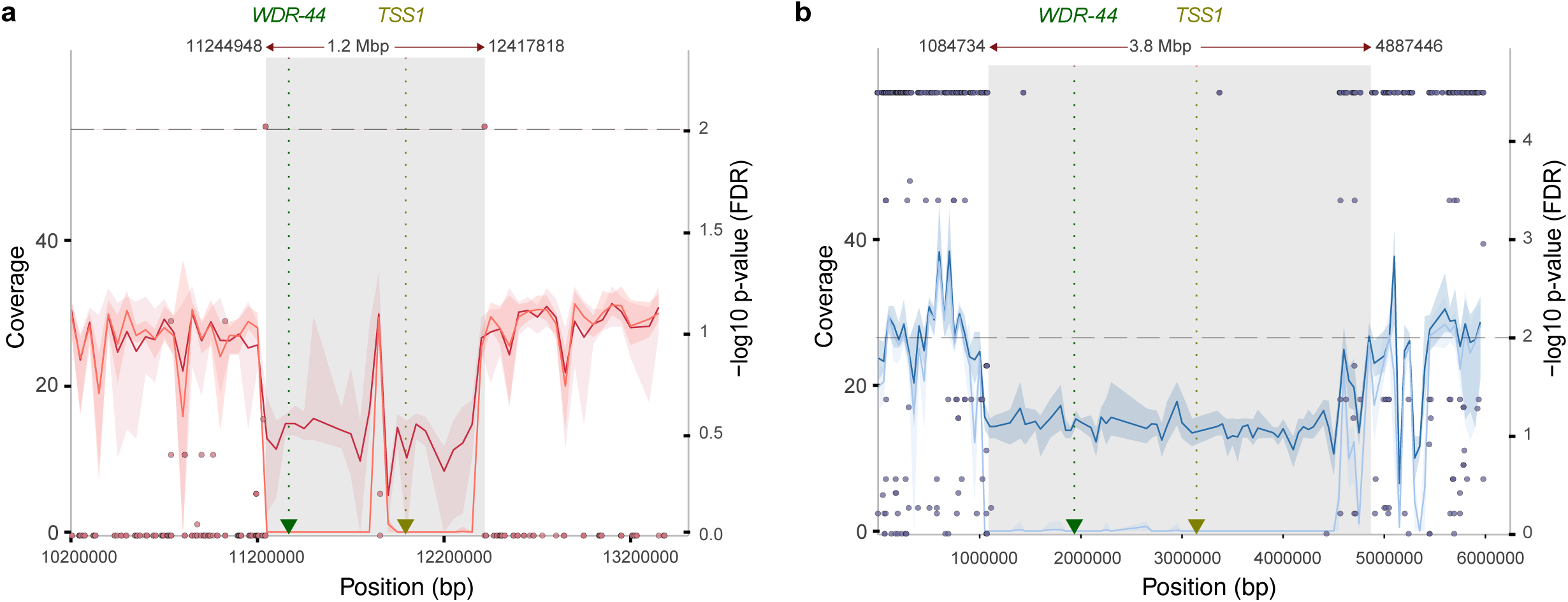
*S-*loci of *Linum grandiflorum* and *Linum perenne* harbor large S-morph hemizygous regions containing distyly candidate genes. Both *L. grandiflorum* contig h1tg000023l (**a**) and *L. perenne* contig ht1g000002l (**b**) harbor S-morph hemizygous *S-*linked regions (coverage values, left y-axis) which contain key candidate distyly genes *TSS1* and *WDR-44*. The size of the hemizygous region and the pattern of SNP association (points showing GWAS significance vs position, right y-axis, significance level α=0.01 indicated by a dashed line) differ between species. In each plot, darker and lighter lines correspond to S-morph and L-morph normalized coverage, respectively, surrounded by shaded regions indicating 95% confidence intervals. The grey areas correspond to regions hemizygous in S-morph individuals, based on coverage analysis and alignment of haplotype-resolved assemblies. The positions of key candidate genes *TSS1* and *WDR-44* are indicated by dotted lines and arrows and the x-axis shows position on each contig (in base pairs).

Finally, inspection of the *L. grandiflorum* genome annotation showed that the S-morph specific region on the dominant *S-*allele (i.e., on h1tg000023l) harbored the *L. grandiflorum* gene *Thrum Style Specific 1* (*TSS1*), a style length candidate gene previously identified as *S-*linked (Ushijima *et al*., 2015) (Figs 2a, S1). Taken together, these results indicate that the *S-*locus of *L. grandiflorum* consists of a ∼1.2 Mb genomic region which harbors presence-absence variation and is hemizygous in S-morph individuals.

In *L. perenne*, family-based GWAS analysis resulted in significant associations between floral morph and SNP genotype on four contigs. We identified a total of 124 significantly associated SNPs on contig ht1g000002l, 33 on contig ht1g000009l, 2 on ht1g000026l, and 20 on ht1g000047l of our hap1 haplotype-resolved genome assembly (Fisher exact test, assuming dominant effect of the S-morph-specific allele, FDR<0.01) (Figs 2b, S1). These results were validated using a population-based GWAS which showed that the same four contigs accounted for 97.7% of GWAS hits (Note S2).

Both contigs with the highest number of morph-associated SNPs (ht1g000002l and ht1g000009l on hap1) map to the same contig in the alternate haplotype-resolved genome assembly (ht2g000035l on hap2), implying that at least 87.7% of the SNPs that show an association with floral morph map to the same chromosome. In total, the contigs showing an association between floral morph and SNP genotype span more than 30 Mb in *L. perenne.* The large size of the region associated with floral morph is likely due to elevated linkage disequilibrium in this specific genomic region (Note S3; Fig. S2).

Coverage analyses further indicate that hap1 contig ht1g000002l corresponds to the dominant *S-*allele, as it harbored an ∼3.8 Mb region specific to the S-morph (median normalized coverage 14.7 in S-morph, 0 in L-morph) (Fig. 2b), and additionally identified an ∼800 kb region specific to the recessive allele (hemizygous in S-morph, diploid in L-morph). Inspection of the annotation of the dominant *S-*haplotype showed the presence of an ortholog of *TSS1* (Figs 2b, S1; Table 1; Data S1B). These results indicate that the *L. perenne S-*locus includes an approximately 3.8 Mb region that is specific to and hemizygous in the S-morph (Figs 2b, S1).

**Table 1.**
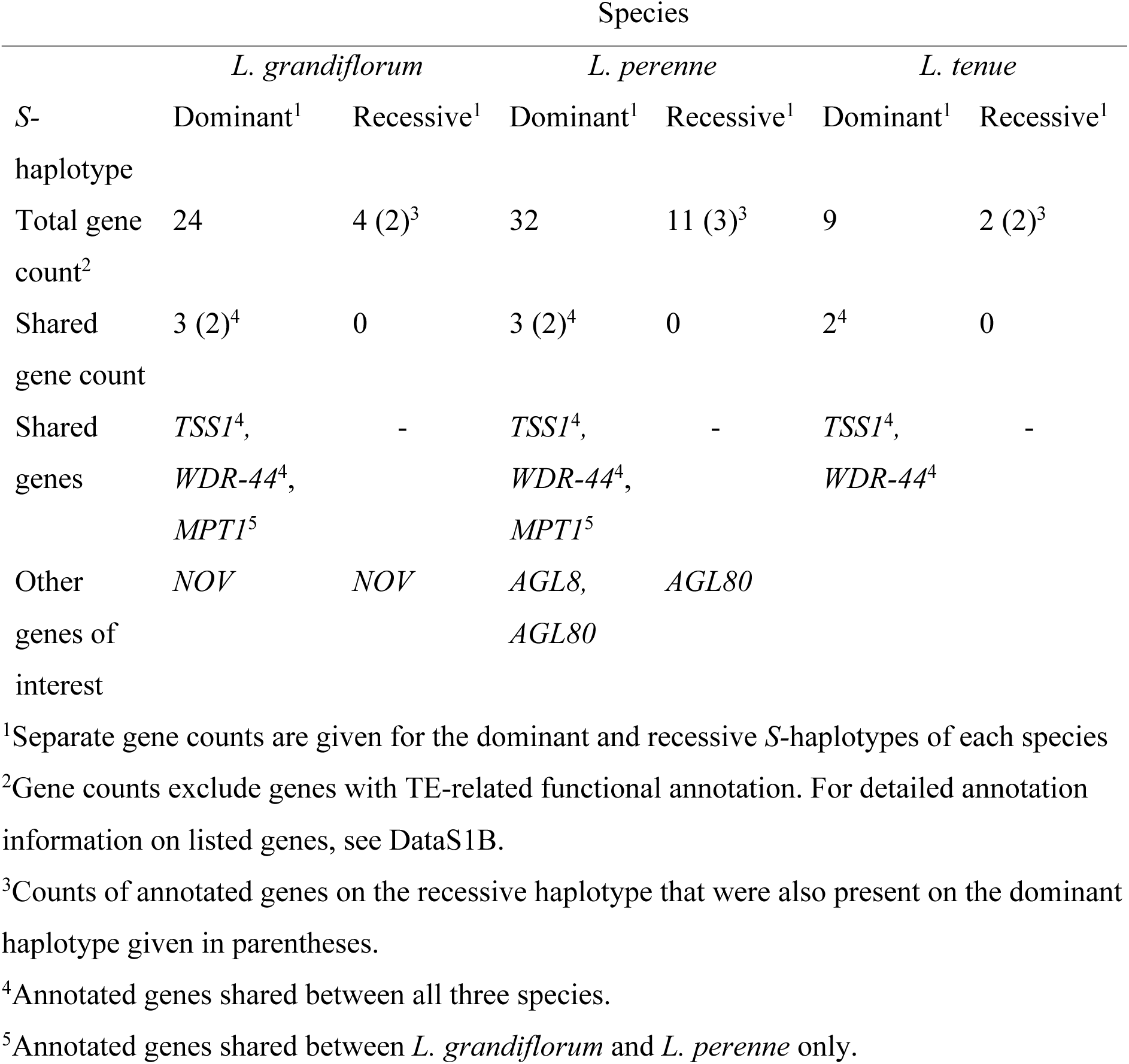
Total number of protein-coding genes annotated at the dominant and recessive *S-*hemizygous region of *Linum grandiflorum*, *Linum perenne*, and *Linum tenue*, the number and identity of genes shared between species, and the number of other genes of potential interest for floral morphology/distyly.

Taken together, association mapping, coverage analyses, and comparisons of haplotype-resolved assemblies indicate that, like in *L. tenue*, the distyly *S-*loci of *L. perenne* and *L. grandiflorum* harbor indel variation, but their S-morph-specific hemizygous regions differ in size (*L. perenne* ∼3.8 Mb, *L. grandiflorum* ∼1.2 Mb). Both *S-*loci are considerably larger than the previously identified *S-*locus of *L. tenue*: 260 kb (Gutiérrez-Valencia et al. 2022).

### Divergent S-loci share distyly candidate genes despite pervasive differences in gene content

Next, we compared the gene content of the *S-*loci of *L. perenne, L. grandiflorum* and *L. tenue*. Apart from the style length candidate gene *TSS1* ((Ushijima *et al*., 2015); also termed *LtTSS1* (Gutiérrez-Valencia *et al*., 2022)), the *S-*loci of both *L. perenne* and *L. grandiflorum* harbored orthologs of the anther height/pollen SI candidate gene *WDR-44* (also termed *LtWDR-44* (Gutiérrez-Valencia *et al*., 2022, 2024)) identified as *S-*linked in *L. tenue* (Gutiérrez-Valencia *et al*., 2022, 2024). One additional *S*-linked gene, *MPT1* (mitochondrial phosphate transporter; GO:0005315) was shared between *L. perenne* and *L. grandiflorum*, but not with the more distantly related *L. tenue* (Fig. 3a-b). No other gene homology was detected when comparing the gene content of the hemizygous region of *L. grandiflorum* to the four morph-associated contigs of *L. perenne* (Fig. 3a-b, S3; Table 1; Note S4). The number of annotated genes in the *S-*linked hemizygous region differed greatly between *L. grandiflorum, L. perenne* and *L.* tenue, with the longer, dominant haplotype having 24 vs 32 annotated protein-coding genes in *L. grandiflorum* and *L. perenne,* compared to only nine in *L. tenue* (Gutiérrez-Valencia et al. 2022) (Table 1, Data S1B).

**Fig. 3.**
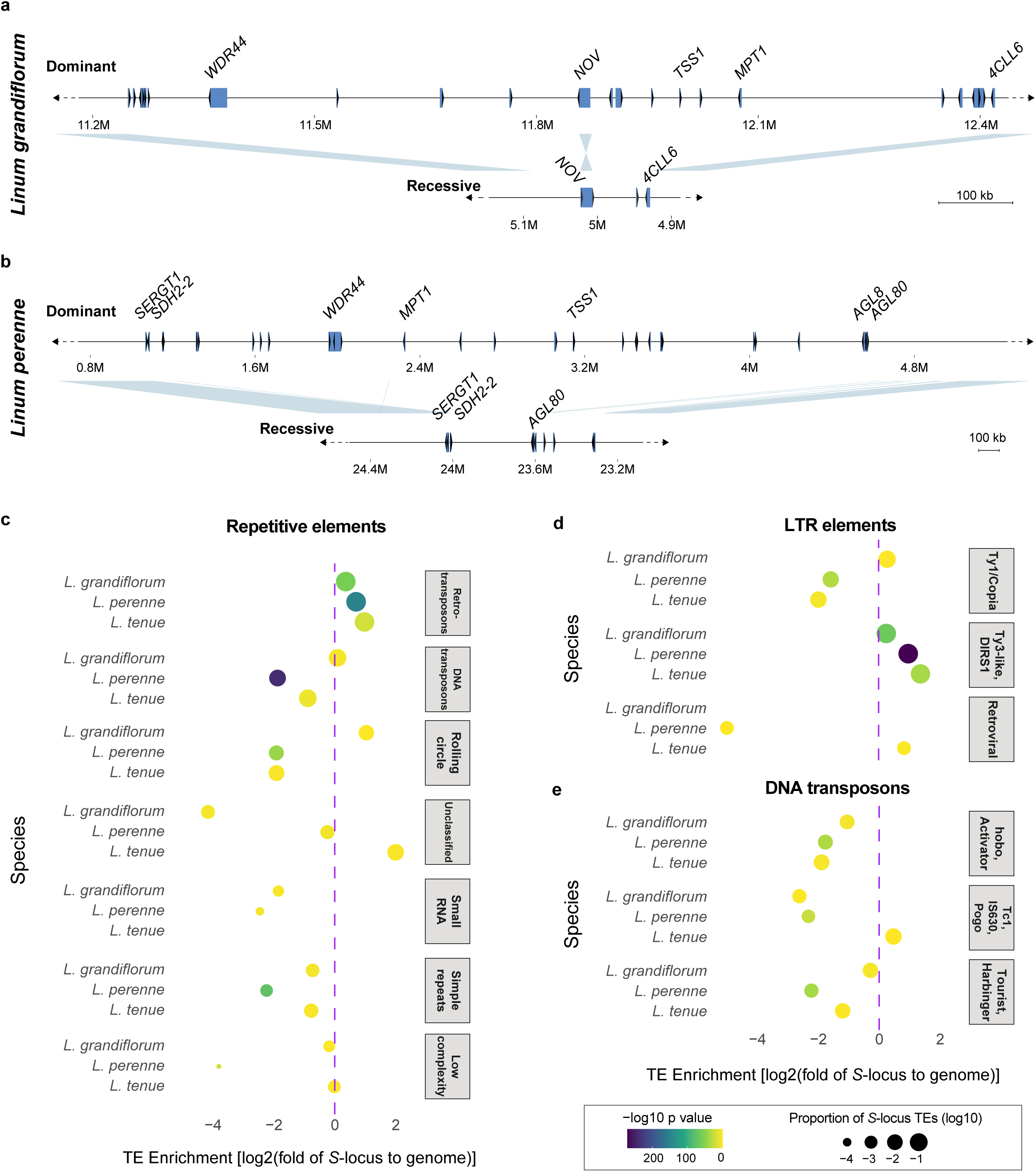
Gene and repetitive element content at the hemizygous *S-*locus regions. **a-b.** Schematic depiction of the haplotype structure and gene content on the dominant and recessive alleles at the hemizygous region of the *L. grandiflorum* (**a**) and *L. perenne* (**b**) *S-* locus. Genes are indicated by blue boxes and arrows indicating orientation. Names of candidate genes (*S-*linked genes shared between *Linum* species) and genes present on both the recessive and dominant haplotypes are shown. **c-e.** The log2-fold enrichment of repetitive elements (**c**), long terminal repeat (LTR) elements (**d**) and DNA transposons (**e**) at the S-morph hemizygous *S-*locus region of *L. grandiflorum*, *L. perenne* and *L. tenue*. Colours indicate the -log10 p-value from a binomial test of repeat enrichment. Circle sizes denote the (log10-transformed) proportion of the *S-*locus region made up of a certain type of repeat.

In *Primula* (Huu *et al*., 2020) and *L. tenue* (Gutiérrez-Valencia *et al*. 2022), the gene set of the *S-*locus has been assembled stepwise via gene duplication. If this process occurred continuously after the origin of the *S-*locus in *Linum*, it might help explain the large differences in gene content between *Linum S-*loci that we observe (Note S4). To test this hypothesis, we estimated *d_S_* between *S-*locus genes and their closest paralogs in *L. grandiflorum* and *L. perenne*, and found evidence for wide differences in the timing of duplication, as well as very recent duplication of *S-*locus genes (Fig. S3). The closest paralogs of *S-*locus genes were found on multiple contigs in both *L. grandiflorum* and *L. perenne*, as expected under stepwise gene duplication (Table S3). These results suggest that stepwise gene duplication, occurring independently in different *Linum* lineages, has contributed to the differences in *S-*locus gene content we observe.

### Divergent S-loci are enriched for different classes of repeats

The *S-*locus is expected to accumulate repeats due to the combined effects of lack of recombination and reduced effective population size (reviewed by Gutiérrez-Valencia et al. 2021). In line with this expectation, we found that the *S-*locus was enriched in repetitive elements relative to the genome-wide average in both *L. perenne, L. grandiflorum,* and *L. tenue* (Fig. 3c). Enrichment was driven primarily by retroelements, specifically Ty3-like Long Terminal Repeat (LTR) retroelements (Fig. 3d). However, the content of certain classes of TEs differed between species, with the *L. grandiflorum S-*locus enriched for rolling circle TEs, in contrast to the *S-*loci of *L. perenne* and *L. tenue* (Fig. 3e).

### Functional constraints on distyly candidate genes at the S-locus over 30 Mya

The presence of *TSS1* and *WDR-44* on the dominant haplotypes of the *S-*loci of *L. grandiflorum, L. perenne* and *L. tenue* (Gutiérrez-Valencia *et al*., 2022) despite major differences in *S-*locus gene content (Note S4) suggests that *TSS1* and *WDR-44* may be ancestrally shared and conserved at the *S-*locus due to their function in the determination of floral morph and/or SI. A simple molecular clock analysis of synonymous divergence at *TSS1* and *WDR-44* between *L. perenne* and *L. tenue* supported this conclusion, as it placed the split between these species at approximately 31-37 Mya (*TSS1: t =* 36.6 Mya (± S.E. 7.3 Mya), *d_S_=*0.513±0.10*; WDR-44: t*=31.3 Mya (± S.E. 2.1 Mya), *d_S_=*0.438±0.03), consistent with these genes having been retained since the diversification of *Linum* ∼33 Mya.

While *TSS1* is a single-copy gene with homologs in outgroups of *Linum*, *WDR-44* is part of a gene family and harbors non-*S-*linked paralogs (Gutiérrez-Valencia *et al*., 2022). To determine when *TSS1* and *WDR-44* first came together in the same genomic region, and to quantify sequence-level constraint on these distyly candidate genes, we assembled sequences of *TSS1, WDR-44* and a set of paralogs of *WDR-44* from five additional distylous *Linum* species (Figs 1a, 4a-b; Table S1). Phylogenetic analysis indicated that the *S*-linked copy of *WDR-44* originated by gene duplication approximately 37 Mya (95% highest posterior density interval (HPD): 30.43-48.22 Mya), suggesting that duplication and translocation of *WDR-44* into a genomic region already harboring *TSS1* occurred at or before the diversification of *Linum* (estimated to have occurred 33 Mya (95% HPD: 27.2-38.3 Mya) (Maguilla *et al*., 2021) (Figs 4b, S4). Consistent with this hypothesis, the closely related outgroup *T. sinensis* only harbored a sequence clustering with the non-*S-*linked paralogs of *WDR-44*, while *WDR-44* sequences of more distant outgroups fell outside of the Linaceae. Taken together, these results support our inference that *WDR-44* duplication occurred at or around the time of diversification of *Linum* (Fig. S4).

**Fig. 4.**
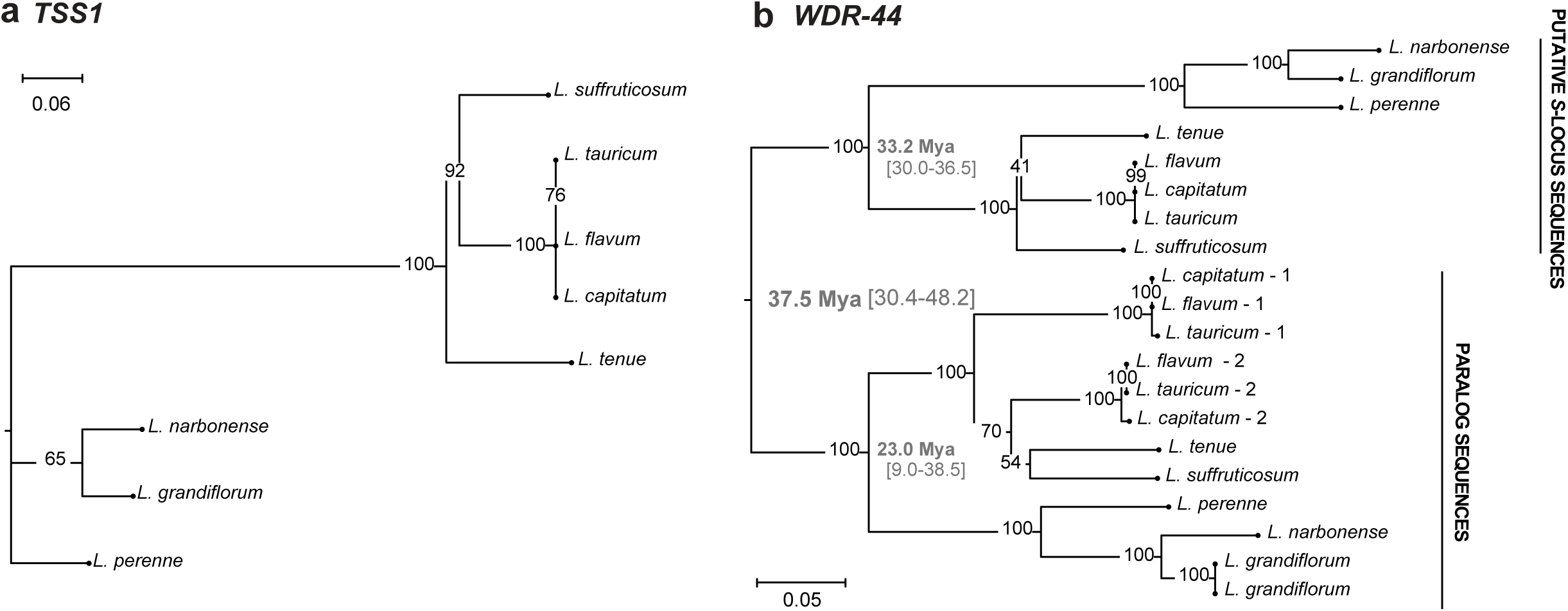
Phylogenetic trees of *TSS1* and *WDR-44* for eight style polymorphic species of *Linum*. **a.** Phylogenetic tree of the conserved *S-*locus candidate genes *TSS1* for style polymorphic *Linum* (all distylous except *L. grandiflorum* which exhibits stigma height dimorphism) reconstructed using RAxML under a GTR-GAMMA substitution model. Support values based on 100 bootstraps are indicated by each node. **b.** Phylogenetic tree of the conserved *S-*locus candidate genes *WDR44* for style length polymorphic *Linum* reconstructed using RAxML under a GTR-GAMMA substitution model. Support values based on 100 bootstraps are indicated by each node. Estimates of the inferred timing of duplication and age of each clade based on BEAST2 analysis are shown for *WDR-44*, with 95% confidence intervals indicated in square brackets.

If *TSS1* and *WDR-44* were retained at the distyly *S-*locus in *Linum* for >30 Mya, we would expect these genes to be under functional constraint. To test this hypothesis, we analyzed ratios of non-synonymous to synonymous divergence (*d_N_/d_S_*) across our eight *Linum* species. We found that for both *TSS1* and the *S-*linked copy of *WDR-44*, a simple model with a single *d_N_/d_S_* across our *Linum* species was supported (*TSS1*: LRT, log LRT test statistic: 1.93, 8 df, *NS*; *WDR-44:* LRT, log LRT test statistic: 1.18, 8 df, *NS*), and both *d_N_/d_S_* estimates were well below 1 (*d_N_/d_S_* of 0.29±0.05 for *TSS1*, and 0.37±0.03 for *WDR-44*), consistent with both genes being under purifying selection (see also Note S5). However, elevated *d_N_/d_S_* of the *S*-locus copy of *WDR-44* compared to its paralogs (0.37±0.03 vs 0.27±0.02; LRT, log LRT test statistic=9.08, 2 df, P=0.0107), suggests the possibility of relaxed purifying selection or alternatively more frequent positive selection on the *S-*locus copy of *WDR-44*. Elevated *d_N_/d_S_* at the *S-*locus copy might be expected under a model where duplication and neofunctionalization contributed to the formation of the distyly *S*-locus.

These results suggest that the distyly *S-*locus of *Linum* formed at or before the diversification of *Linum*, and that the two *S*-locus candidate genes *TSS1* and *WDR-44,* which are shared among widely diverged distylous *Linum* species, are under purifying selection, possibly related to their role in determining floral morph differences and/or SI.

### Regulation of style length by brassinosteroids in widely divergent distylous Linum

The style length candidate gene *TSS1* was present and conserved at the *S-*loci of *L. tenue*, *L. perenne* and *L. grandiflorum*, all of which exhibit style length polymorphism. We previously hypothesized that *TSS1*, which is primarily expressed in styles of S-morph individuals (Ushijima *et al*., 2015; Gutiérrez-Valencia *et al*., 2022), might result in shorter style cells and thereby shorter styles by downregulating brassinosteroid-responsive genes in a manner similar to its Arabidopsis homolog *VUP1* (Grienenberger & Douglas, 2014). If so, treating floral buds with brassinosteroids should result in longer styles and style cells specifically in S-morph but not in L-morph *Linum* individuals. If the mechanism of action of *TSS1* has remained conserved, we expect the effect of brassinosteroid treatment to be present in widely divergent distylous *Linum* species, as long as their *S-*locus harbors functional *TSS1.* To test this hypothesis, we conducted a brassinosteroid supplementation experiment where L- and S-morph flower buds of *L. tenue* and *L. perenne* were treated with brassinosteroid solution (eBL: 10 μM 24-epibrassinolide, dissolved in 0.1% of the solvent dimethylsulfoxide, DMSO) or control treatment (control: 0.1% DMSO only), followed by measurement of style and stamen length.

In line with our expectation, eBL treatment resulted in significantly longer styles in both *L. perenne* (Table 2; Fig. 5a) and *L. tenue* (Table 2; Fig. 5b) due to the specific effect of eBL treatment on style length in S-morph individuals (*L. perenne*: Two-way ANOVA, interaction of treatment and morph, Table 2; Fig. 5c; *L. tenue*: Two-way ANOVA, interaction of treatment and morph, Table 2; Fig. 5d). On average, eBL treatment resulted in 0.82 mm (95% CI: 0.34-1.30 mm; Fig. 5f) longer styles in *L. perenne* S-morph individuals and 0.94 mm (95% CI: 0.50-1.38 mm) longer styles in *L. tenue* S-morph individuals. While styles were significantly longer in eBL-treated S-morph individuals, they were still shorter than those of L-morph individuals (Fig. 5a), possibly due to the timing of application and/or concentration of eBL treatment used. The eBL treatment had no effect on style length in L-morph individuals of *L. perenne* or *L. tenue* (Fig. 5a, b), and there was no significant interaction effect on stamen length in *L. perenne* or *L. tenue* (Table S4).

**Fig. 5.**
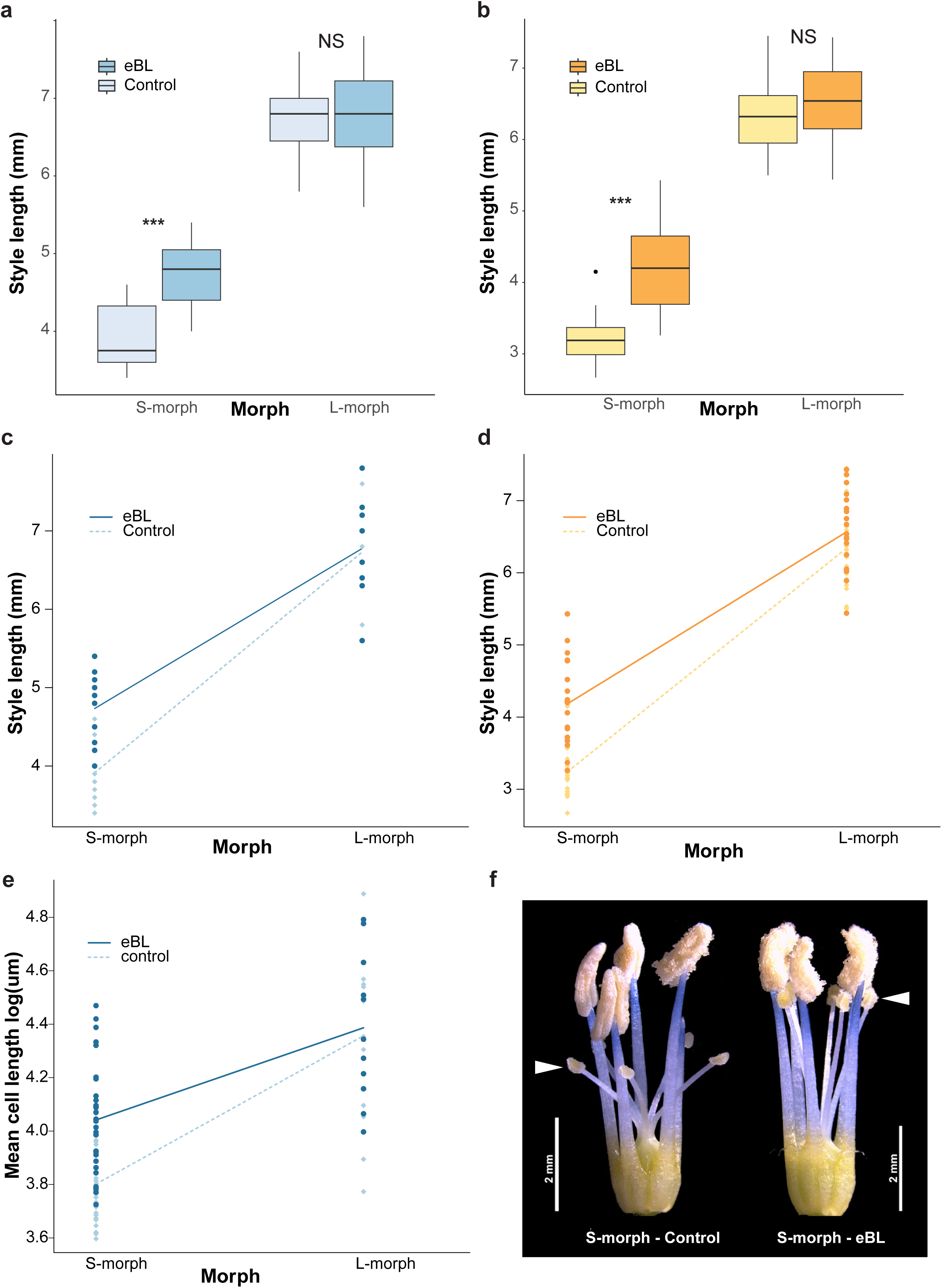
Brassinosteroid supplementation results in longer styles and style cells in S-morph but not L-morph morph individuals of widely divergent *Linum* species. **a, b.** Boxplots showing significantly longer S-morph but not L-morph styles in *L. perenne* (**a**) and *L. tenue* (**b**) after epibrassinolide (10 uM eBL in 0.1% DMSO) treatment of flower buds compared to control treatment (0.1% DMSO only). **c, d.** Interaction plots demonstrating a significant interaction between floral morph and eBL treatment, in both *L. perenne* (**c**) and *L. tenue* (**d**). **e.** eBL treatment results in significantly longer epidermal style cells in S-morph but not L-morph individuals of *L. perenne*. **f.** Photograph of control and eBL-treated *L. perenne* S-morph sexual organs, showing the effect of eBL treatment on style length. Stigma positions are indicated by arrows.

**Table 2.** Brassinosteroid treatment has a morph-specific effect on style length (mm) in both the distylous species *Linum perenne* and *Linum tenue*.

| **Species** | **Source of variation<sup>1</sup>** | **Df<sup>2</sup>** | **SS<sup>3</sup>** | **MS<sup>4</sup>** | **F<sup>5</sup>** | **P-value** |
| --- | --- | --- | --- | --- | --- | --- |
| *L. perenne* | Morph | 1 | 60.36 | 60.36 | 244.24 | <0.0001 |
|  | Treatment | 1 | 3.71 | 3.71 | 15.03 | <0.0001 |
|  | Morph*Treatment | 1 | 1.51 | 1.51 | 6.11 | 0.02 |
|  | Residuals | 42 | 10.38 | 0.25 |  |  |
| *L. tenue* | Morph | 1 | 142.7 | 142.7 | 537.1 | <0.0001 |
|  | Treatment | 1 | 6.48 | 6.48 | 24.4 | <0.0001 |
|  | Morph*Treatment | 1 | 2.44 | 2.44 | 9.2 | 0.003 |
|  | Residuals | 72 | 19.13 | 0.27 |  |  |

<sup>1</sup>Analysis of variation (ANOVA) sources of variation
<sup>2</sup>Degrees of freedom
<sup>3</sup>Sums of squares
<sup>4</sup>Mean squares
<sup>5</sup>F-statistic

To test whether the effect of brassinosteroid treatment on style length was mediated by style cell length, we measured epidermal style cell length in *L. perenne* after eBL and control treatment. There was a significant effect of eBL treatment on mean style cell length (*F_1,75_, P*<0.0001; Figs 5e, S5a; Table S5), as well as a significant interaction between eBL treatment and morph (F_1,75_=5.7, P=0.02; Fig. 5e; Table S5). The effect of eBL on mean style cell length in the S-morph was 12.6 μm (95% CI: 4.9-20.3 μm). The brassinosteroid treatment had no significant effect on mean style cell length in L-morph individuals (Fig. 5e; Fig. S5).

Taken together, the impact of brassinosteroid treatment on style and style cell length specifically in S-morph individuals suggests that a mechanism relying on the brassinosteroid pathway, likely regulated by *TSS1*, contributes to style length differences between floral morphs in widely diverged distylous *Linum*.

## Discussion

One of the most prominent examples of convergent floral evolution in plants is distyly, yet until recently little was known about the molecular nature of distyly *S-*loci and the mechanisms underlying this multi-trait balanced polymorphism. Here, we expand our understanding of the nature and evolution of distyly *S*-loci by leveraging haplotype-resolved genome assemblies of widely diverged *Linum* species.

We show that the *S*-locus supergenes of three *Linum* species that diverged as far back as 33 Mya share the same genetic architecture, characterized by an S-morph specific hemizygous region, although the sizes of the hemizygous regions vary greatly from ∼260 kb to ∼3.8 Mb. All three *Linum* species further harbor two only shared genes at the S-morph specific region of their *S-*loci; the style length candidate gene *TSS1* (Ushijima *et al*., 2015; Gutiérrez-Valencia *et al*., 2022) and anther position/pollen SI candidate gene *WDR-44* (Gutiérrez-Valencia *et al*., 2022, 2024). The presence of *WDR-44* at the *S-*locus of *L. grandiflorum*, which lacks stamen length variation, is especially intriguing, given that *WDR-44* has been hypothesized to be an anther height/pollen SI gene (Gutiérrez-Valencia *et al*., 2022). We cannot currently rule out conservation of *WDR-44* in *L. grandiflorum* due to an effect of this gene on pollen SI, as *L. grandiflorum* shares heteromorphic SI with both *L. tenue* and *L. perenne* (Murray 1986). Further detailed characterization and functional work will be required to determine the effects of *WDR-44* in *Linum*.

We have previously shown that *TSS1* is present in outgroups of *Linum* (Gutiérrez-Valencia *et al*., 2022), and here we use paralog dating to show that *WDR-44* originated through gene duplication ∼37 Mya (95% CI 30.4 Mya - 48.2 Mya), suggesting that these two distyly candidate genes became colocated in one genomic region at or before the diversification of *Linum* s.l. ∼33 Mya (Maguilla *et al*., 2021). The distyly *S-*locus therefore probably evolved early during diversification of *Linum*, through a process involving duplication and likely also neofunctionalization of *WDR-44*. Duplication and neofunctionalization of the anther height gene *GLO2* (*GLO^T^*) have previously been documented at the distyly supergene in *Primula* (Li *et al.,* 2016; Huu *et al*., 2020).

Our results suggest that *TSS1* was present and could have evolved presence-absence polymorphism regulating style length before *WDR-44* was duplicated and became co-located with *TSS1*, broadly in line with predictions of the “pollen transfer” model of the evolution of distyly (Lloyd & Webb, 1992). However, we cannot currently rule out other scenarios, including one where both stamen and style length polymorphism was established at the same time through a large indel generating presence-absence variation for both *TSS1* and *WDR-44*. Therefore, while our results do not allow us to distinguish between major models for the evolution of distyly (Charlesworth & Charlesworth, 1979; Lloyd & Webb, 1992), they suggest the distyly *S-*locus evolved early during diversification of *Linum*.

Molecular evolutionary analyses of *TSS1* and *WDR-44* show that both of these *S-*locus genes are under purifying selection in *Linum*. To test whether sequence conservation is related to effects on distyly, we focused on *TSS1*, which has been hypothesized to regulate style length via its impact on the brassinosteroid pathway (Gutiérrez-Valencia *et al*., 2022a). While functional studies of *TSS1* are required to validate our findings, morph-specific effects of exogenous brassinosteroid application on style and style cell length in *L. perenne* suggest that *TSS1* governs style length through its effect on brassinosteroid-regulated genes. Our finding that brassinosteroid supplementation also had a morph-specific effect on style length in *L. tenue* implies that the mechanism underlying style length polymorphism is conserved across these widely diverged distylous *Linum* species. Future studies should investigate whether brassinosteroid treatment also affects female SI reaction in *Linum*. While we cannot fully rule out the involvement of additional hormonal pathways in the regulation of style length in *Linum*, our results suggest that genes impacting the brassinosteroid pathway have repeatedly been recruited during convergent evolution of style length polymorphism in distylous species, including in *Primula* (Huu *et al*., 2022) and *Turnera* (Matzke *et al*., 2020, 2021).

An early cessation of recombination at the supergene followed by independent evolution for an extended period could possibly explain the extensive variation in size, gene and repeat content that we observe among *Linum* distyly supergenes. Indeed, structural variants and repeats are expected to accumulate in non-recombining regions, especially over long evolutionary timescales (reviewed in (Gutiérrez-Valencia *et al*., 2021b)). Our observations of differences in the accumulation of repeat classes at the distyly *S-*locus among the studied species are consistent with independent TE accumulation in different lineages. Recent genomic studies have documented gene movement into the *S-*locus (termed “stepwise assembly”; (Huu *et al*., 2020; Gutiérrez-Valencia *et al*., 2022)). In *L. tenue*, we previously found evidence for very recent gene movement into the *S-*locus (Gutiérrez-Valencia *et al*., 2022), suggesting that this process could still be ongoing. Our analyses here suggest that stepwise gene duplication, occurring independently and continuously in different *Linum* lineages, could help explain the marked differences in gene content we observe at *Linum S*-loci.

Our study revealed strong differences in the gene content at the *S-*locus across divergent *Linum* species, despite the presence of conserved candidate genes and a conserved hormone-dependent mechanism regulating style length. Similar findings have been reported in other systems that share features with distyly supergenes. For instance, distyly supergenes and sex determining regions share the features of recombination suppression between alleles and morph-specific inheritance of one allele (reviewed in Gutiérrez-Valencia *et al*. 2021b). Therefore, it is perhaps not surprising that many of the patterns we document at the *Linum* distyly supergene are reminiscent of those at the sex determining region of *Actinidia* species (Akagi *et al*., 2023). Specifically, while we observe only two shared candidate genes for distyly located in an S-morph-specific genomic region across divergent *Linum* species, Akagi et al., (2023) similarly observed only three shared candidate sex determination genes located in a male-specific genomic region in divergent *Actinidia* species. Like Akagi et al., (2023), we also find that molecular mechanisms underlying morphs appear conserved, despite marked sequence-level evolution of the morph-specific region. These similarities suggest that further investigation of the parallels between the evolution of plant mating system supergenes and sex determining regions is a fruitful avenue for future theoretical and empirical work.

Taken together, our results shed light on the genetic architecture, origin and evolution of the *Linum* distyly supergene, revealing the presence of conserved candidate genes and pathways regulating distyly, despite marked differences in supergene size, gene and repeat content. Our results and the genome assemblies produced here provide a foundation for further work on the role of parallel genetic changes for convergent evolution of floral form and function in distylous species.

## Acknowledgements

We thank Benjamin Laenen and Aurélie Desamoré for assistance with plant sampling, Alireza Foorozani, P. William Hughes, and Juanita Gutiérrez-Valencia for assistance with plant cultivation and flow cytometry, Jerker Eriksson for technical assistance with plant growth chambers, Tomas Larsson for bioinformatics advice and Matias Wanntorp for bioinformatic assistance. This project has received funding from the European Research Council (ERC) under the European Union’s Horizon 2020 and Horizon Europe research and innovation programmes (grant agreement No 757451 and 101132305), from the Swedish Research Council (grant agreements 2019-04452 and 2023-04532), from the Erik Philip-Sörensen foundation to T.S., and from the Nilsson-Ehle foundation to P.I.Z. Z.P. was funded by a Carl Tryggers foundation grant (CTS21:1471) to T.S. The authors acknowledge support from the National Genomics Infrastructure (NGI) in Sweden, funded by Science for Life Laboratory, the Knut and Alice Wallenberg Foundation and the Swedish Research Council. Long-read sequencing was performed at the NGI Uppsala Genome Center, whereas short-read sequencing was performed by the NGI SNP&SEQ Technology Platform in Uppsala. Hi-C sequencing was performed by NGI in Stockholm. The computations were enabled by resources in projects SNIC2022/22-683, SNIC 2022/22-695, NAISS 2023/22-129, NAISS 2024/5-158, and NAISS 2023/4-5 provided by the National Academic Infrastructure for Supercomputing in Sweden (NAISS) at UPPMAX, funded by the Swedish Research Council through grant agreement no. 2022-06725. Support by the National Bioinformatics Infrastructure Sweden is gratefully acknowledged.

## Author Contributions

T.S. conceived of and designed the study, acquired funding, supervised the work and wrote the original draft. A.L. performed experiments. P.I.Z., Z.P., M.F., L.S., E.P.W., I.B., A.C. and T.S. performed analyses. P.I.Z., Z.P. and A.L. revised and edited the manuscript, with additional comments and input from M.F., L.S., E.P.W., I.B., and A.C. P.I.Z. and Z.P. contributed equally.

## Data Availability

All sequencing data generated in this study has been uploaded to the European Nucleotide Archive (ENA, https://www.ebi.ac.uk/ena/) and will be released upon acceptance. Original code has been uploaded to Zenodo (10.5281/zenodo.14500764). NBIS open-source pipelines for genome annotation are available at: https://github.com/NBISweden/GAAS; https://github.com/NBISweden/AGAT; https://github.com/NBISweden/pipelines-nextflow.

