## Supporting Information for "Genomic studies in *Linum* shed light on the evolution of the distyly supergene and the molecular basis of convergent floral evolution"

The following Supporting Information is available for this article:

**Fig. S1** *S*-loci of *Linum grandiflorum* and *Linum perenne* harbor large S-morph hemizygous regions containing key distyly candidate genes.
**Fig. S2** Extended linkage disequilibrium in natural populations of *Linum perenne* along *S*-locus contig h1tg000002l.
**Fig. S3** Synonymous divergence between *S*-locus genes and their closest paralogs in *Linum grandiflorum* and *Linum perenne*.
**Fig. S4** Phylogenetic tree of *WDR-44*.
**Fig. S5** Brassinosteroid treatment results in longer style cells in S-morph individuals of *Linum perenne*.
**Table S1** Origin and section classification of plant material used in this study.
**Table S2** Genome assembly statistics for Hifiasm Hi-C integrated haplotype-resolved assemblies.
**Table S3** Synonymous divergence for all genes with paralogs at the dominant allele *S*-hemizygous regions and their closest paralogs, in *Linum grandiflorum* and *Linum perenne*.
**Table S4** Brassinosteroid treatment has no detectable morph-specific effect on stamen length (mm) in *Linum perenne* or *Linum tenue*.
**Table S5** Analysis of log-transformed mean cell length data from the epibrassinolide supplementation experiment shows that brassinosteroid treatment has a morph-specific effect on style cell length (mm) in *Linum perenne*.
**Note S1.** Assembly contamination screening and masking results.
**Note S2.** Population-based validation of GWAS results.
**Note S3.** Extensive linkage disequilibrium at the *L. perenne* distyly S-locus.
**Note S4.** Description of gene content at the S-loci of *L. grandiflorum* and *L. perenne*.
**Note S5.** Comparative molecular evolutionary analysis of rates of evolution of *TSS1* and *WDR-44.*

**Fig. S1 *S*-loci of *L. grandiflorum* and *L. perenne* harbor large S-morph hemizygous regions containing key distyly candidate genes.** Both *L. grandiflorum* (**a-b**) and *L. perenne* (**c-d**) harbor S-morph hemizygous *S*-linked regions (coverage values, left y-axis) which contain key candidate distyly genes *TSS1* and *WDR-44*. The size of the hemizygous region and the pattern of SNP association (points showing GWAS significance vs position, right y-axis, significance level α=0.01 indicated by a dashed line) differ between species, with morph-associated SNPs found across the entire *S*-locus contig h1tg000002l in *L. perenne*. In each plot, darker and lighter lines correspond to S-morph and L-morph depth of coverage, respectively, surrounded by shaded regions indicating 95% confidence intervals. The grey areas correspond to regions hemizygous in S-morph individuals, based on coverage analysis and alignment of haplotype-resolved assemblies. The positions of candidate genes *TSS1* and *WDR-44* are indicated by dotted lines and arrows.

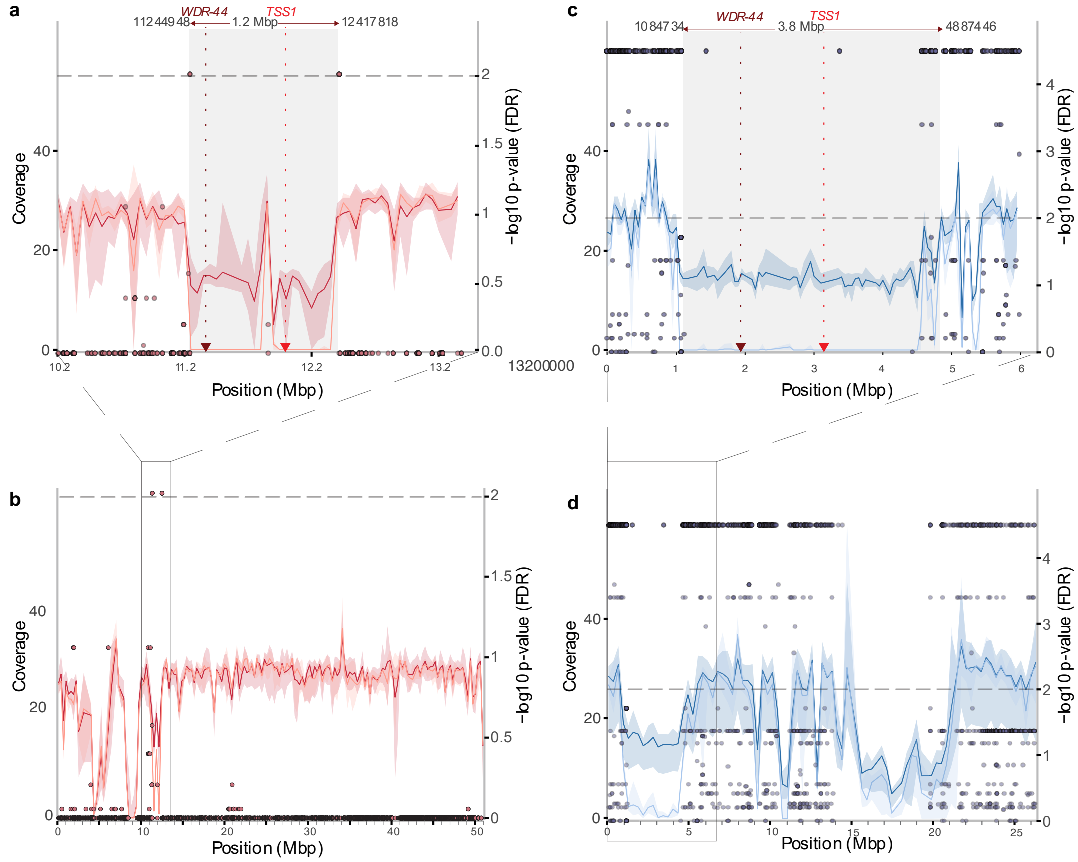

**Fig. S2 Extended linkage disequilibrium in natural populations of *L. perenne* along *S*-locus contig h1tg000002l.** Heatmaps of median $r^2$ values between all pairs of windows of 100 kb along the *S*-locus contig ht1g000002l and the control contig ht1g00031l for three natural populations of *L. perenne* (panels a-c). White areas in heatmap represent lack of SNPs. The lower plots show SNP number per window along each contig. LD distributions along the two contigs were significantly different for the two populations (Wilcoxon rank sum test, ger3: W = 355251530, p-value<0.001, ger5: W = 631400752, p-value<0.001, ger2: W = 600311350, p-value<0.001).

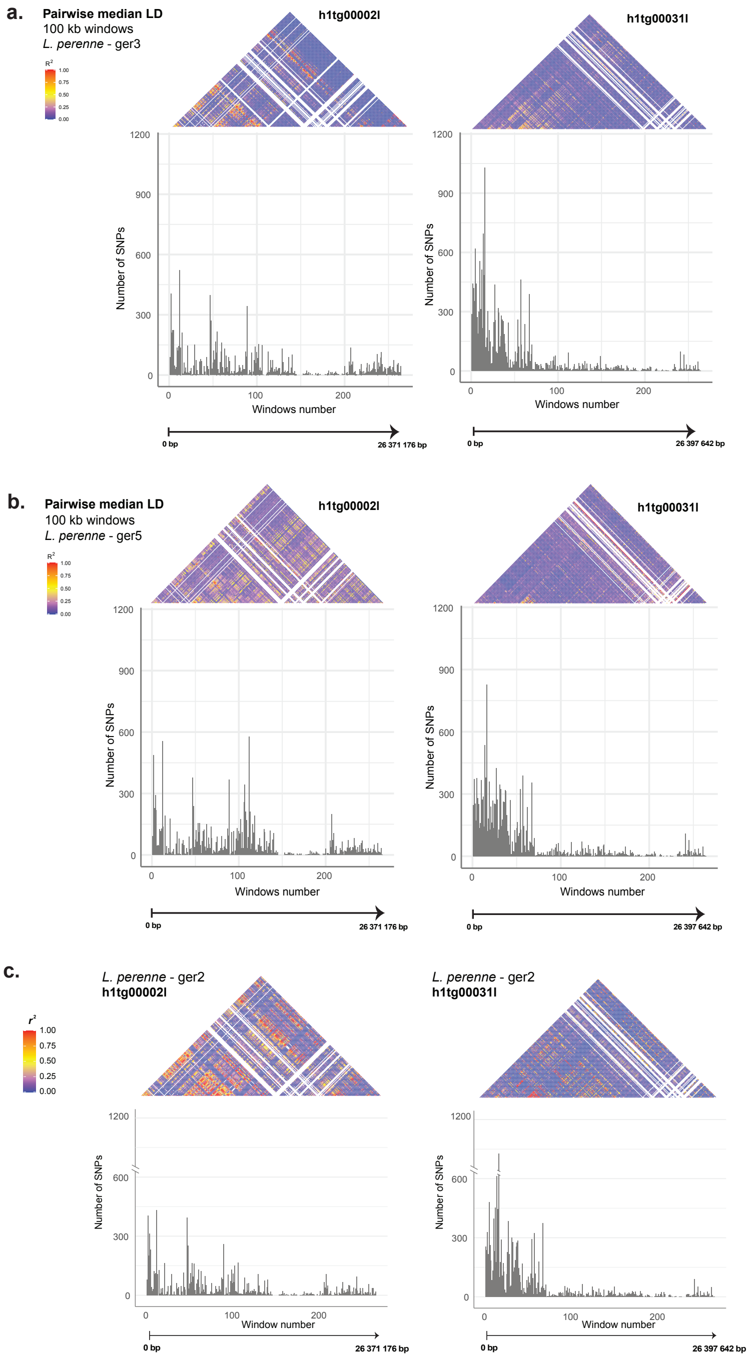

**Fig. S3 Synonymous divergence between *S*-locus genes and their closest paralogs in *L. perenne* and *L. grandiflorum*.** The solid line represents the divergence time between *WDR-44* and its paralogs, while the dashed lines indicate the 95% confidence interval for this divergence time. Variation in synonymous divergence values between the different *S*-locus genes and their paralogs across both species supports a stepwise formation of the gene set at the *S*-locus. Additionally, for some genes, the synonymous divergence is lower than the estimated emergence time of the *S*-locus in *Linum*, suggesting that these genes were incorporated into the *S*-locus after its initial formation.

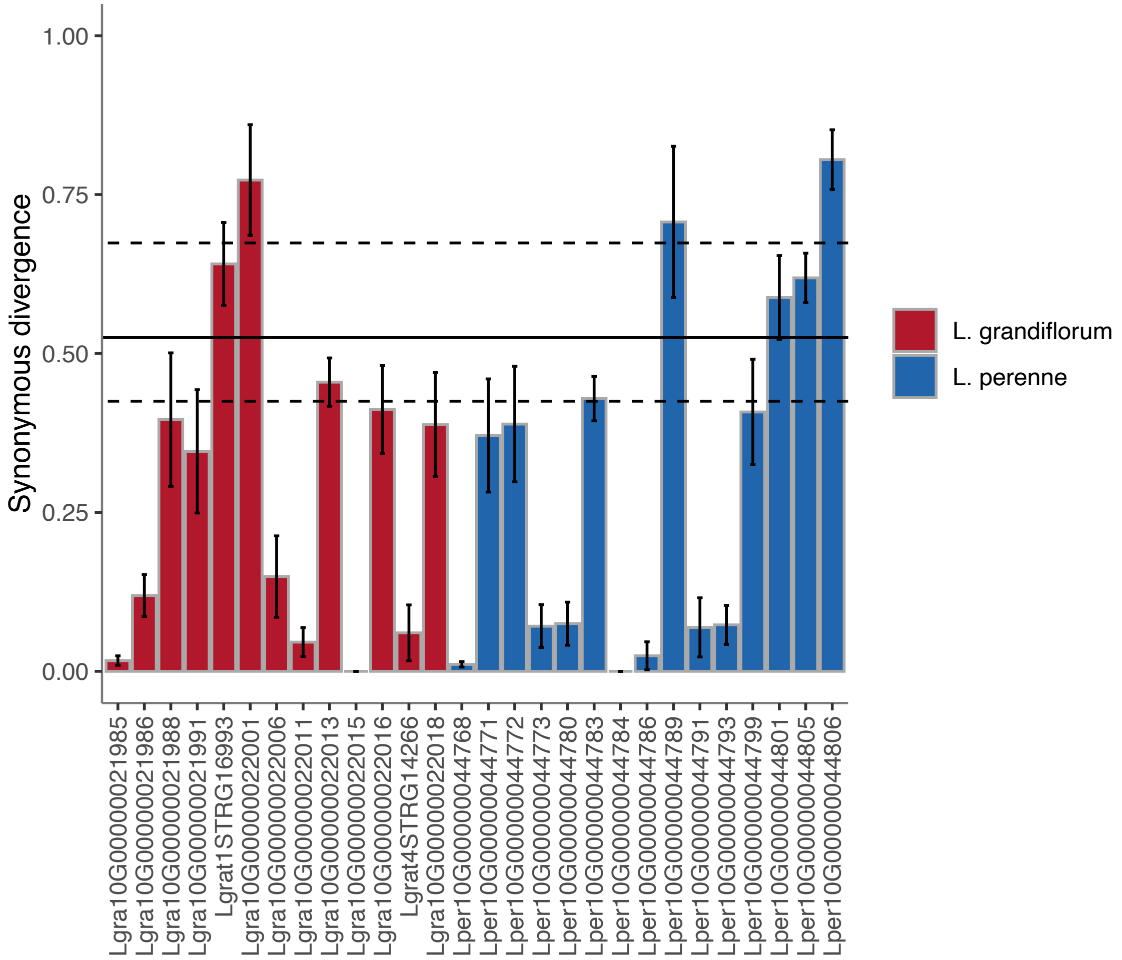

**Fig. S4 Phylogenetic tree of *WDR-44*.** This tree was reconstructed with RAxML and contains *S*-linked *WDR-44* sequences, paralog sequences and the sequence for one closely related outgroup (*Tirpitzia sinensis*) and two more distant outgroups (*Manihot esculenta* and *Populus trichocarpa*, respectively). The *Tirpitzia* sequence falls within the paralog cluster, supporting the emergence of the *S*-locus copy at the time of diversification of *Linum*.

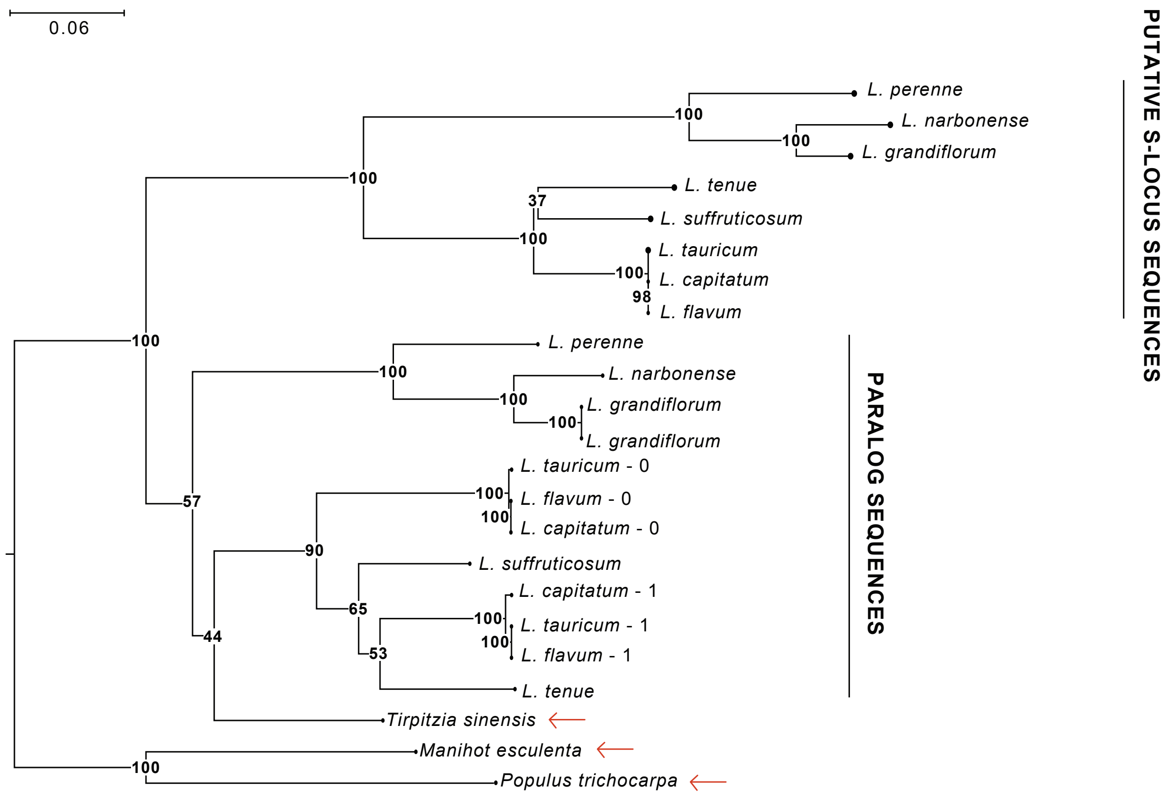

**Fig. S5 Brassinosteroid treatment results in longer style cells in S-morph individuals of *L. perenne*. a.** Effects of epibrassinolide (eBL) treatment on mean style cell length (μm) across all style sections. Mean style cell length was significantly longer after eBL treatment in S-morph (thrum) but not L-morph (pin) individuals. *** indicates P-value<0.001. NS indicates not significant. **b-d.** Boxplots showing mean style cell length with eBL or control treatment separately for each style section. **e.** Photograph indicating the three style sections where cells were measured, separately for an S-morph plant (left) and an L-morph plant (right). **f.** Example of micrograph of style cells after control (left) and eBL (right) treatment of an S-morph (thrum) plant.

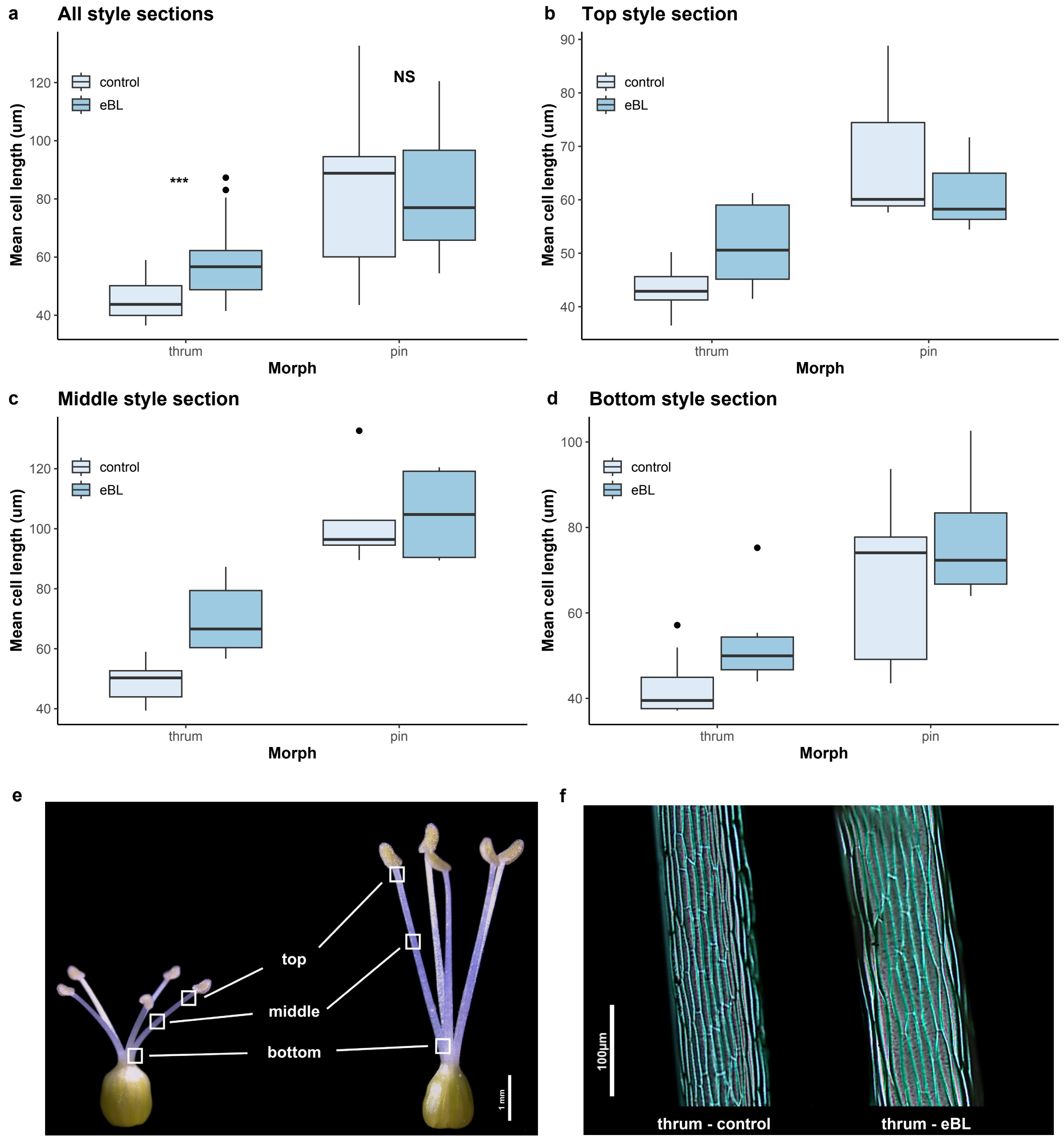

**Table S1 Origin and section classification of plant material used in this study.**

| Species | Accession/origin | Section |
|---|---|---|
| *Linum capitatum* | LIN 1903 (IPK) | Syllinum |
| *Linum flavum* | LIN 2001 (IPK) | Syllinum |
| *Linum grandiflorum* | LIN 1389 (IPK) | Linum |
| *Linum grandiflorum* | LIN 974 (IPK) | Linum |
| *Linum grandiflorum* | LIN 10 (IPK) | Linum |
| *Linum narbonense* | LIN 1807 (IPK) | Linum |
| *Linum perenne* | LIN 2003 (IPK) | Linum |
| *Linum perenne* | ger2 near Othfresen, Germany | Linum |
| *Linum perenne* | ger3 near Marienhagen, Germany | Linum |
| *Linum perenne* | ger5 near Goslar, Germany | Linum |
| *Linum suffruticosum* | 18-5-2 near Ronda, Spain | Linopsis C |
| *Linum tauricum* | LIN 1658 (IPK) | Syllinum |
| *Linum tenue* | CL, near Castillo de Locubín, Spain | Linopsis B |
| *Linum tenue* | SMT, near Santa María de Trassiera, Spain | Linopsis B |

[1] Accessions labeled IPK are derived from the IPK Gatersleben Gene Bank and the IPK accession code is given in Accession/origin.

**Table S2 Genome assembly statistics for Hifiasm Hi-C integrated haplotype-resolved assemblies.**

| | ***L. perenne*** **assemblies** | | | ***L. grandiflorum*** **assemblies** | | |
|---|---|---|---|---|---|---|
| **Statistic** | **hap1** | **hap2** | **primary** | **hap1** | **hap2** | **primary** |
| **No. of contigs** | 905 | 514 | 809 | 856 | 436 | 647 |
| **Total length[1]** | 739,225,655 | 753,366,752 | 773,825,471 | 816,779,520 | 743,719,982 | 809,872,110 |
| **Largest contig** | 41,207,707 | 49,950,927 | 87,425,665 | 90,933,564 | 60,021,352 | 125,429,925 |
| **GC (%)** | 41.45 | 41.67 | 41.46 | 42 | 41.81 | 41.86 |
| **N50[2]** | 10,271,661 | 11,313,697 | 23,398,707 | 28,074,137 | 22,188,264 | 69,915,768 |
| **N75[3]** | 4,808,757 | 5,584,177 | 14,389,690 | 15,858,823 | 12,991,397 | 38,093,872 |
| **L50[4]** | 19 | 17 | 10 | 9 | 12 | 5 |
| **L75[5]** | 44 | 41 | 21 | 18 | 22 | 8 |
| **BUSCO** | 95.1 | 94.8 | 94.7 | 94.4 | 94 | 94.4 |

[1]Flow-cytometry based genome size estimates for *L. perenne* and *L. grandiflorum* are 784 vs 908 Mb.

[2]N50: 50% of contigs in the assembly are longer than the N50 length.

[3]N75: 75% of the contigs in the assembly are longer than the N75 length.

[4]L50 Count of the smallest number of contigs which together make up 50% of the genome assembly size.

[5]L75: Count of the smallest number of contigs which together make up 75% of the genome assembly size.

**Table S3 Synonymous divergence for all genes with paralogs at the dominant allele *S*-hemizygous regions and their closest paralogs in *Linum grandiflorum* and *Linum perenne*.**

| ***S*-locus gene** | **Paralog** | **Contig** | **Start** | **End** | $d_S$[1] | **SE**[2] |
|---|---|---|---|---|---|---|
| Lgra10G00000021985 | Lgra10G00000008224 | h1tg000005l | 31463522 | 31465015 | 0.0168 | 0.0074 |
| Lgra10G00000021986 | Lgra10G00000031222 | h1tg000019l | 6180865 | 6181227 | 0.119 | 0.033 |
| Lgra10G00000021988 | Lgra10G00000028065 | h1tg000010l | 2169113 | 2170729 | 0.396 | 0.105 |
| Lgra10G00000021991 | Lgra10G00000026991 | h1tg000006l | 29657130 | 29658098 | 0.346 | 0.097 |
| t1STRG16993 | Lgra10G00000018032 | h1tg000014l | 11676715 | 11686899 | 0.641 | 0.065 |
| Lgra10G00000022001 | Lgra10G00000009030 | h1tg000005l | 72849596 | 72850303 | 0.773 | 0.087 |
| Lgra10G00000022006 | Lgra10G00000017843 | h1tg000014l | 4161762 | 4167805 | 0.149 | 0.064 |
| Lgra10G00000022011 | Lgra10G00000019904 | h1tg000018l | 20747387 | 20748495 | 0.0459 | 0.0228 |
| Lgra10G00000022013 | Lgra10G00000024773 | h1tg000016l | 4164938 | 4167333 | 0.455 | 0.038 |
| Lgra10G00000022015 | Lgra10G00000018894 | h1tg000021l | 17187215 | 17188497 | 0.000 | 0.000 |
| Lgra10G00000022016 | Lgra10G00000017915 | h1tg000014l | 7437038 | 7440734 | 0.412 | 0.069 |
| t4STRG14266 | Lgra10G00000036514 | h1tg000009l | 21362014 | 21366348 | 0.0604 | 0.0440 |
| Lgra10G00000022018 | Lgra10G00000019815 | h1tg000018l | 18975568 | 18975744 | 0.388 | 0.082 |
| Lper10G00000044768 | Lper10G00000015926 | h1tg000818l | 27001 | 32440 | 0.0109 | 0.0042 |
| Lper10G00000044771 | Lper10G00000041146 | h1tg000102l | 1490252 | 1493370 | 0.371 | 0.089 |
| Lper10G00000044772 | Lper10G00000028692 | h1tg000071l | 2098032 | 2105393 | 0.389 | 0.091 |
| Lper10G00000044773 | agat-gene-1 | h1tg000002l | 1586118 | 1586688 | 0.0710 | 0.0337 |
| Lper10G00000044780 | Lper10G00000015230 | h1tg000052l | 5534072 | 5534446 | 0.0749 | 0.0339 |
| Lper10G00000044783 | Lper10G00000025512 | h1tg000059l | 1943219 | 1945331 | 0.429 | 0.035 |
| Lper10G00000044784 | Lper10G00000016519 | h1tg000021l | 35147299 | 35147439 | 0.000 | 0.000 |
| Lper10G00000044786 | Lper10G00000040280 | h1tg000046l | 6310639 | 6313807 | 0.0243 | 0.0220 |
| Lper10G00000044789 | Lper10G00000036747 | h1tg000065l | 9284490 | 9284729 | 0.707 | 0.119 |
| Lper10G00000044791 | Lper10G00000034992 | h1tg000003l | 1586489 | 1590067 | 0.0690 | 0.0465 |
| Lper10G00000044793 | Lper10G00000044869 | h1tg000002l | 7081115 | 7082187 | 0.0730 | 0.0307 |
| Lper10G00000044799 | Lper10G00000004039 | h1tg000051l | 13488572 | 13489794 | 0.408 | 0.083 |
| Lper10G00000044801 | Lper10G00000041292 | h1tg000004l | 448836 | 449093 | 0.588 | 0.066 |
| Lper10G00000044805 | Lper10G00000003205 | h1tg000051l | 3100528 | 3101385 | 0.619 | 0.039 |
| Lper10G00000044806 | Lper10G00000008228 | h1tg000025l | 2083158 | 2083955 | 0.805 | 0.047 |

[1]Synonymous divergence

[2]Standard error

**Table S4 Brassinosteroid treatment has no detectable morph-specific effect on stamen length (mm) in *Linum perenne* or *Linum tenue*.**

| Species | Source of variation[1] | Df[2] | SS[3] | MS[4] | F[5] | *P* |
|---|---|---|---|---|---|---|
| *L. perenne* | Morph | 1 | 31.23 | 32.23 | 54.09 | <0.0001 |
| | Treatment | 1 | 0.59 | 0.59 | 1.02 | 0.32 (NS) |
| | Morph*Treatment | 1 | 0.32 | 0.32 | 0.55 | 0.46 (NS) |
| | Residuals | 40 | 23.09 | 0.58 | | |
| *L. tenue* | Morph | 1 | 113.3 | 113.3 | 417.4 | <0.0001 |
| | Treatment | 1 | 1.22 | 1.22 | 4.50 | 0.04 |
| | Morph*Treatment | 1 | 0.009 | 0.009 | 0.0334 | 0.86 (NS) |
| | Residuals | 72 | 19.55 | 0.27 | | |

[1]Analysis of variance sources of variation
[2]Degrees of freedom
[3]Sums of squares
[4]Mean squares
[5]F-statistic

**Table S5 Analysis of log-transformed mean cell length data from the epibrassinolide supplementation experiment shows that brassinosteroid treatment has a morph-specific effect on style cell length (mm) in *Linum perenne*.**

| Source of variation[1] | Df[2] | SS[3] | MS[4] | F[5] | P-value |
|---|---|---|---|---|---|
| Morph | 1 | 3.47 | 3.47 | 121.8 | <0.0001 |
| Segment | 2 | 1.44 | 0.72 | 25.3 | <0.0001 |
| Treatment | 1 | 0.66 | 0.66 | 23.3 | <0.0001 |
| Morph*Segment | 2 | 0.21 | 0.11 | 3.7 | 0.03 |
| Morph*Treatment | 1 | 0.16 | 0.16 | 5.7 | 0.02 |
| Residuals | 75 | 2.13 | 0.03 | | |

[1]Analysis of variance sources of variation
[2]Degrees of freedom
[3]Sums of squares
[4]Mean squares
[5]F-statistic

### Supplementary Notes

**Note S1 Assembly contamination screening and masking results.**
In *L. perenne* two contigs (h1tg000166c and h1tg000316l) were removed due to high coverage (>200x) and similarity to plastid or mitochondrial DNA, while four regions of 88.4 kb, 59 kb, 186.4 kb, and 78.6 kb were masked in contigs h1tg000014l, h1tg000021l, h1tg000036l, and h1tg000092l, respectively, due to signs of contamination. In *L. grandiflorum*, two contigs (h1tg000046c and h1tg000187c) were removed due to high similarity to plastid or mitochondrial DNA.

**Note S2 Population-based validation of GWAS results.**
Because family-based GWAS analysis has limitations in terms of resolution, we conducted a follow-up validation analysis based on 53 samples from a natural population. Analyzing a total of 10,681 SNPs in 53 individuals from one natural population of *L. perenne* (population ger3), we identified a total of 220 SNPs on six contigs that were significantly associated with floral morph (FDR<0.05). Only five SNPs mapped to contigs not identified in the family-based analysis, and the remaining 215 SNPs  (97.7%) mapped to the same contigs as in the family-based analysis). The two major contigs identified in the family-based analysis harbored 85.5% of the SNPs significantly associated with floral morph, and the length of the associated genomic regions amounted to 32.1 Mb. These results support those of our family-based analysis.

**Note S3 Extensive linkage disequilibrium at the *L. perenne* distyly S-locus.**
To test for limited recombination in a large genomic region around the *S*-hemizygous region in *L. perenne*, we estimated linkage disequilibrium (LD) across the ~26 Mb *S*-locus contig (h1tg000002l). For this purpose, we analyzed population genomic data from three natural populations of *L. perenne* from Germany (ger2, n=63; ger3, n=53; ger5, n=20; Table S1). . We assessed LD by estimating $r^2$ between each pair of SNPs along the *S*-locus contig as well as for a control contig of similar size (h1tg000031l) in PLINK v1.90b4.9 (Purcell *et al.*, 2007). We aggregated $r^2$ values in bins of 100 kb and plotted LD heatmaps with these bins for both contigs using *Ldheatmap* v1.0-5 (Shin *et al.*, 2006) in R v.4.4.0. We compared $r^2$ values for the two contigs using a Wilcoxon rank sum test in R v.4.4.0.

We found elevated LD along the length of the *S*-locus contig, whereas that was not the case for the control contig (Fig. S2), suggesting that the extended LD is not a result of non-equilibrium demographic processes. The extended LD at the *S*-locus in natural populations of *L. perenne* suggests that the genomic region with limited recombination is more extensive in *L. perenne* than in *L. grandiflorum*, where GWAS hits were in the immediate vicinity of the S-hemizygous region (Fig. 2a-b). Likewise, analyses of LD around the *L. tenue S*-locus found no evidence of extended LD outside of the S-hemizygous region (Gutiérrez-Valencia *et al.*, 2022).

**Note S4 Description of gene content at the S-loci of *L. grandiflorum and L. perenne*.**
The number of annotated genes in the *S*-linked hemizygous region differed greatly between *L. tenue*, *L. grandiflorum* and *L. perenne,* with the longer, dominant haplotype having only nine protein-coding genes in *L. tenue* (Gutiérrez-Valencia *et al.*, 2022), in contrast to those of *L.*

*grandiflorum* and *L. perenne,* which harbored 24 and 32 protein-coding genes, respectively. In *L. grandiflorum*, the thrum-hemizygous region harbored nine genes with functional annotation, including *TSS1*, *WDR-44*, *MPT1*, one gene involved in auxin-mediated developmental responses (*NOV1*) and five additional genes with various functions not clearly related to floral development or reproduction (Fig. 4a; Data S1B). Two of the genes present on the dominant *S*-haplotype (*NOV* and *4CLL6*) were also present on the recessive *S*-haplotype. The remaining 15 protein-coding genes were of unknown function and five of them overlapped to more than 50% with annotated repeats (Fig. 4a; Data S1B).

In *L. perenne* the longer, dominant *S*-haplotype harbored 12 genes with functional annotation, including *TSS1*, *WDR-44*, *MPT1*, as well as two genes with functions related to floral or reproductive development: two MADS-box transcription factors (*AGL8* and *AGL80*) involved in floral meristem development and female gametophyte cell differentiation, respectively (Fig. 4b; Data S1B). The remaining functionally annotated genes had no clear connection to floral morphology or reproduction, and 14 out of 20 genes without functional annotation showed more than 50% overlap with annotated repeats (Data S1B).

**Note S5 Comparative molecular evolutionary analysis of rates of evolution of TSS1 and WDR-44.**

We tested whether rates of evolution ($d_N/d_S$) at *TSS1* and *WDR-44* were elevated in relation to those of other genes across the genomes of *L. grandiflorum, L. perenne* and *L. tenue*. To do so, we compared estimates of $d_N/d_S$ at *TSS1* and *WDR-44* in *L. grandiflorum, L. perenne* and *L. tenue* to those at a set of 3,864 1-1-1 orthologs, identified using OrthoFinder v2.5.5 (Emms and Kelly, 2019). We aligned the orthologs with PRANK (Löytynoja 2013) and we found no evidence for more extreme $d_N/d_S$ values at either *TSS1* or *WDR-44* compared to 1-1-1 orthologs, consistent with purifying selection (mean $d_N/d_S$ for 1-1-1 orthologs: 0.18, 95% CI 0.05-0.41, for *TSS1*: 0.32, for *WDR-44*: 0.35).
